# Pan-active imidazolopiperazine antimalarials target the *Plasmodium falciparum* intracellular secretory pathway

**DOI:** 10.1101/735894

**Authors:** Gregory M. LaMonte, Danushka S. Marapana, Nina Gnadig, Sabine Ottilie, Madeline R. Luth, Tilla S. Worgall, Frances Rocamora, Gregory M. Goldgof, Roxanne Mohunlal, T.R Santha Kumar, Jenny K. Thompson, Edgar Vigil, Jennifer Yang, Dylan Hutson, Trevor Johnson, Jianbo Huang, Roy M. Williams, Bing Yu Zou, Andrea L. Cheung, Prianka Kumar, Timothy J. Egan, Marcus C.S. Lee, Dionicio Siegel, Alan F. Cowman, David A. Fidock, Elizabeth A. Winzeler

**Affiliations:** Department of Pediatrics, University of California, San Diego, School of Medicine, La Jolla, CA 92093, United States; Division of Infection and Immunity, Walter and Eliza Hall Institute for Medical Research; Department of Microbiology & Immunology, Columbia University Irving Medical Center, New York, New York 10032, USA; Department of Pathology and Cell Biology, Columbia University Irving Medical Center, New York, New York 10032, USA; Department of Laboratory Medicine, University of California, San Francisco; Department of Chemistry, University of Cape Town, Rondebosch, South Africa; Skaggs School of Pharmacy and Pharmaceutical Sciences, University of California, San Diego, School of Medicine, La Jolla, CA 92093, USA; Parasites and Microbes Programme, Wellcome Sanger Institute, Hinxton. CB10, 1SA, UK; Division of Infectious Diseases, Department of Medicine, Columbia University Irving Medical Center, New York, New York 10032, USA

## Abstract

One of the most promising new compound classes in clinical development for the treatment of malaria is the imidazolopiperazines (IZPs) class. Human trials have demonstrated that members of the IZP series, which includes KAF156 (Ganaplacide) and GNF179, are potent and effective against *Plasmodium* symptomatic asexual blood-stage infections. Unlike other commonly used antimalarials, they also prevent transmission and block future infection in animal models. Despite the identification of several *Plasmodium falciparum* resistance mechanisms including mutations in ER-localized PfCARL (PfEMP65), Acetyl-coA transporter, and PfUGT transporter, IZP’s mechanism of action remains unknown.

To investigate, we combined *in vitro* evolution and whole-genome analysis in the model organism *Saccharomyces cerevisiae* with molecular, metabolomic, and chemogenomic methods, in *P. falciparum*. *S. cerevisiae* clones that resist IZP activity carry multiple mutations in genes that encode endoplasmic reticulum(ER)-based lipid homeostasis and autophagy including *elo2*, *elo3*, *sur2*, *atg15* and *lcb4*, as well as ER-based *sec66.* In *Plasmodium*, IZPs cause inhibition of protein trafficking, block the establishment of new permeation pathways and result in ER expansion. We also observe sensitization with other secretion inhibitors such as brefeldin A and golgicidin as well as synthetic lethality with PfSEC62. Our data show that IZPs target the secretory pathway and highlight a novel mechanism for blocking parasite growth and development that is distinct from those of standard compounds used to treat malaria. In addition, we provide physiological signatures and hallmarks for inhibitors that work through this mechanism of action and show that IZPs are tool compounds for studying ER-dependent protein processing in different species.

## Introduction

Malaria remains the most common human parasitic disease, sickening an estimated 219 million people annually and causing an estimated 435,000 deaths worldwide^1^. Although a reduction in the malaria burden has been achieved through the combined use of public health infrastructure, vector control and increased access to medical treatment^2,3^, these efforts are hindered by a variety of factors, such as the lack of an effective vaccine, increased mosquito resistance to pesticides, and the continuing emergence of drug-resistant parasite strains ^4–6^. There are also now reports of resistance to nearly all current classes of antimalarials^7^. Given that an estimated 3.2 billion people live in endemic regions and are at risk for malaria,^3^ new classes of antimalarial compounds will be required to combat, and ultimately eradicate this disease.

One of the most promising classes of compounds are the imidazolopiperazines (IZPs), such as KAF156^8^ and its closely related analog GNF179^9^, which differs from KAF156 by a single halogen substitution (F to Cl). The chemotype that led to KAF156 was first identified in a phenotypic screen designed to discover compounds with activity against parasite asexual blood stages^10^. The compound series attracted more attention when it was noted that multiple analogs were present in the library, which all showed activity in a liver stage model that predicts causal antimalarial prophylaxis^9^. Subsequent rounds of medical chemistry^11,12^ led to the development of KAF156, an orally available, simple-to-synthesize compound suitable for testing in humans^8^.

KAF156 shows potent activity against asexual blood (IC_50_ ~ 6 nM), hepatic (IC_50_ = 4.5 nM) and sexual stages (IC_50_ = 5 nM)^8^, and is considered safe with good pharmacokinetic properties in healthy human volunteers^13^. Because the compound was able to dramatically reduce parasite numbers in patients with *P. vivax* and *P. falciparum* patients in a single agent trial^14^ it has now progressed to phase IIb clinical trials (<u>NCT03167242</u>) where it is being tested in combination with lumefantrine. It also has impressive prophylactic activity: treating mice with a single 10 mg/kg oral dose fully protects them from mosquito-borne malaria infection. Furthermore, IZPs have gametocytocidal activity and including them in a blood meal or pretreating with IZPS prevents parasites from being transmissible to mosquitoes^8^, both *in vitro* and *in vivo*. If licensed, IZPs could be clinically superior to current gold standard treatments, such as artemisinin-based combination therapies, which often just provide symptomatic relief and neither prevent malaria from developing nor prevent its transmission. KAF156 could serve as a potent tool in the mission to eliminate this disease.

Previous studies using *in vitro* evolution and whole-genome analysis in *P. falciparum* parasites showed that resistance to IZPs is mediated by mutations in three different genes, *pfcarl*, the cyclic amine resistance transporter (PF3D7_0321900)^8,9^; *pfact*, the Acetyl-CoA transporter (PF3D7_1036800); and *pfugt*, the UDP-galactose transporter (PF3D7_1113300)^15^. GFP-tagging and localization experiments have shown that these three transmembrane transporters are all localized to the endoplasmic reticulum (ER)/Golgi apparatus in *Plasmodium*. Multiple resistance-conferring alleles have been recovered independently in *pfact* and *pfcarl*.

It is unlikely that the nonessential *pfact* encodes the target, although in human cells the *pfact* ortholog, AT-1, appears essential^15^: Parasite mutants with stop codons as well as frameshifts are readily recovered after KAF156 treatment, although these may render the parasites less fit. Like the parasite protein, the human protein is also localized to the ER where it serves to import Acetyl-CoA for use in lysine acetylation of some newly synthesized protein. Its disruption in human cells results in a proteasome-independent ERAD(II) mechanism involving the unfolded protein response and autophagy of the ER^15^. In humans, mutating lysines for some proteins such as BACE results in proteins that are retained in aggregates in the ER. Many *Plasmodium* proteins are often acetylated^16^, often at conserved residues, although it is not clear that this happens in the ER and which acetyl transferases are responsible for this.

*Pfugt* encodes a member of the Solute Carrier 35 Family. Members of this family play a role in import of sugars to the ER/Golgi where most glycoconjugate synthesis occurs^17^. Disruption of some orthologs in worms and plants also lead to ER stress^18,19^. Although disruption mutants in *pfugt* have not been obtained in high-throughput approaches in *P. falciparum*, *pfugt* is a small gene and could have been missed for random reasons^20^. Although PfCARL appears essential^20^, mutations in *pfcarl* confer resistance to unrelated compounds^21,22^ and resistance-conferring mutations in *pfcarl* are located in transmembrane regions and not in an obvious catalytic site. PfCARL, although conserved in evolution, remains understudied, but its yeast ortholog, Emp65 (Endoplasmic Reticulum Membrane Protein 65) protects folding polypeptides from promiscuous degradation^23^. Mutations in all three parasite proteins may lead to slower rates of protein folding, processing and sorting.

Parasites treated with IZPs have also been subjected to metabolic profiling along with other clinical compounds with known modes of action. Allman *et al.*^24^ measured changes in 113 metabolites after treatment with KAF156. These data did not show a clear metabolic perturbation, in contrast to inhibitors of cytochrome bc1, dihydroorotate dehydrogenase, PfATP4, or dihydrofolate reductase, many of which are also active in both blood and hepatic stages.

Given the clinical potential of GNF179, determining its mechanism of action could reveal important new druggable pathways, suggest synergistic drugs that could be used in combination therapies, and provide clues on possible toxicity. Here we report on a series of experiments in *P. falciparum* and *S. cerevisiae* to discern the mode of action of this important antimalarial compound series.

## Results

### Identifying potential targets of GNF179 using *S. cerevisiae* model system genetics

For target identification, we first used *in vitro* evolution studies in the model system *Saccharomyces cerevisiae*, a strategy that has been used to discover the target of cladosporin, a tRNA synthetase inhibitor that acts against both *P. falciparum* and *S. cerevisiae*. IZPs are moderately active against an attenuated strain of *Saccharomyces* that had been genetically modified by replacing 16 ABC multi-drug transporter genes with modified *Aequorea victoria* GFP (eGFP) and that has been dubbed the “Green Monster” ^25^. Altogether 13 different, independent IZP-resistant yeast lines were created by growing the cells for a minimum of 20 generations in the presence of increasing GNF179 concentrations until resistance was observed (with a minimum 1.5x IC_50_ increase for GNF179). Clonal lines were isolated from each resistant culture and retested for sensitivity. The observed resistant strains exhibited 1.5 – 3.1-fold resistance relative to the drug-naïve Green Monster strain.

We sequenced drug-resistant genomes to an average 47.3x coverage and compared these to the drug-sensitive parent (Table S1). Excluding mutations in repetitive elements, we detected 67 total variants including 49 missense, frameshift or nonsense variants, 2 synonymous variants, 2 inframe deletions and 14 intergenics (Table S2). The high proportion of coding to silent mutations suggests that most changes have a beneficial effect. Multiple independent alleles were found in six different genes arising in independent selections (*sec66*(2 variants), *elo2*(5), *lcb4*(3), YMR102C(4), *atg15*(4) and *sur2*(3)), with the most resistant lines harboring multiple mutations (e.g. GNF179-R19g2 with YMR102C, *atg15* and *sur2*) although not generally in catalytic domains (Figure 1a). With ~5800 genes in the genome, the probability of finding more than one allele by chance in the same gene after *in vitro* evolution is very low (*p* = 4.4 e −7) and highlights the reproducibility of the selections. In addition, the set of 38 variants included singletons in the closely related genes *sur1*, *elo3*, and *atg22*. This mutational pattern is specific to IZPs and has not been observed for *in vitro* evolution with other antimalarial compounds whose targets have been identified using the Green Monster system^26,27^.

**Figure 1.**
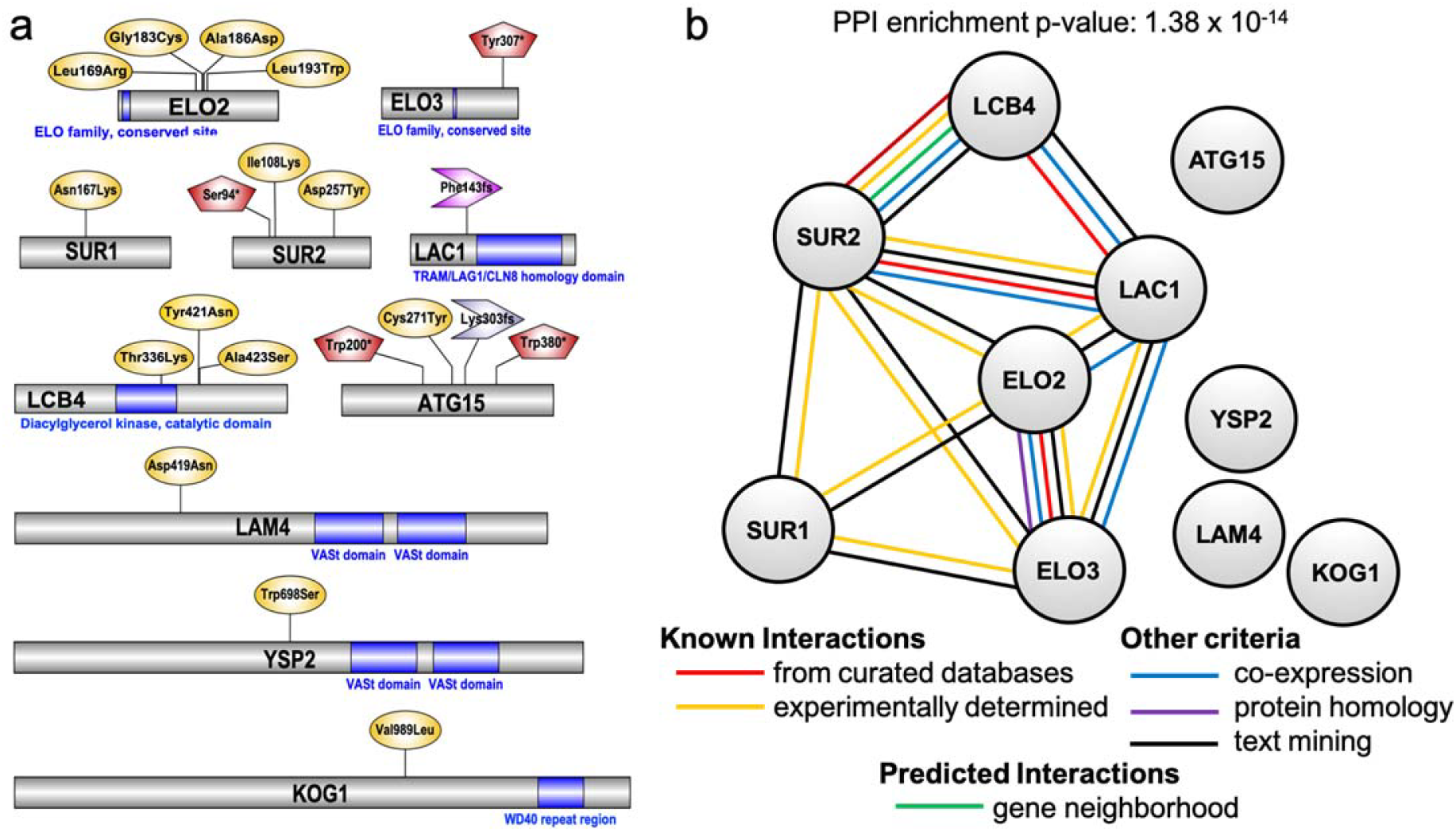
GNF179-resistant yeast strains harbor mutations in endoplasmic reticulum(ER)-based lipid homeostasis and autophagy. (a) Protein maps showing relevant mutations and PROSITE predicted protein domains, if applicable. Maps were generated using Illustrator of Biological Sequences (IBS) software package.^81^ Missense mutations are shown in yellow ovals, nonsense mutations in red pentagons, and frameshift mutations as purple arrow. (b) Protein-Protein Interaction (PPI) network generated using the STRING database.^82^ Each node represents a *S. cerevisiae* protein and connecting lines delineate interactions. The PPI enrichment p-value (p = 1.38 × 10^−14^) indicates that the proteins show significantly more interactions among themselves than would be expected from a random subset of genes from the yeast genome.

With the exception of YMR102C, whose function is not known, these genes are all directly or indirectly associated with trafficking and processes in the ER (Figure 1b). *SEC66* encodes a non-essential subunit of the SEC63 complex that forms a channel competent for SRP-dependent and post-translational SRP-independent protein targeting and import into the ER. *SEC66* disruption is well known to slow the process of protein trafficking, such that trafficking intermediates become evident by gel electrophoresis^28^. Similarly, *ELO2 and ELO3*, which encode fatty acid elongases that contribute to sphingolipid biosynthesis in the ER, were identified in a protein trafficking screen: Alleles in both *ELO2* and *ELO3*, named *VBM1* and *VBM2*,^29^ were identified as suppressors of a v-SNARE mutant in which yeast cells accumulate post-Golgi secretory vesicles, and are defective in invertase secretion.^30^ *ATG15* and *ATG22* play a role in autophagy, which is induced in proteasome-independent ER expansion that results from aggregates of misfolded proteins. *SUR1* (the catalytic subunit of mannosylinositol phosphorylceramide (MIPC) synthase that is required for biosynthesis of mature sphingolipids), *SUR2* (sphingosine hydroxylase involved in sphingolipid metabolism), and *LCB4* (sphingoid long-chain base kinase) are all part of the sphingolipid metabolism pathway, which is carried out in the ER. Sphingolipids and long chain fatty acids play a role in regulating autophagy. The identification of resistance-conferring genes whose products are localized to the ER or Golgi is similar to observations in *P. falciparum*.

As these resistant strains often harbored multiple mutations, we used CRISPR-*Cas9* based genome editing to introduce these mutations in a drug-naïve Green Monster genetic background. For *sec66* (M1I, IC_50_ = 70.1 µM, S107*, IC_50_ = 74.82 µM), *elo2* (G183C IC_50_ = 88.15 µM) and *elo3* (Y307*, IC_50_ = 76.56 µM) vs wild-type (WT) Green Monster IC_50_ = 47.3 µM (Table 2), the CRISPR-edited lines exhibited similar, 1.5 to 1.9-fold levels of resistance to the drug-pressure derived lines, indicating that these three genes were responsible for the resistance observed in those yeast strains.

**Table 1.**
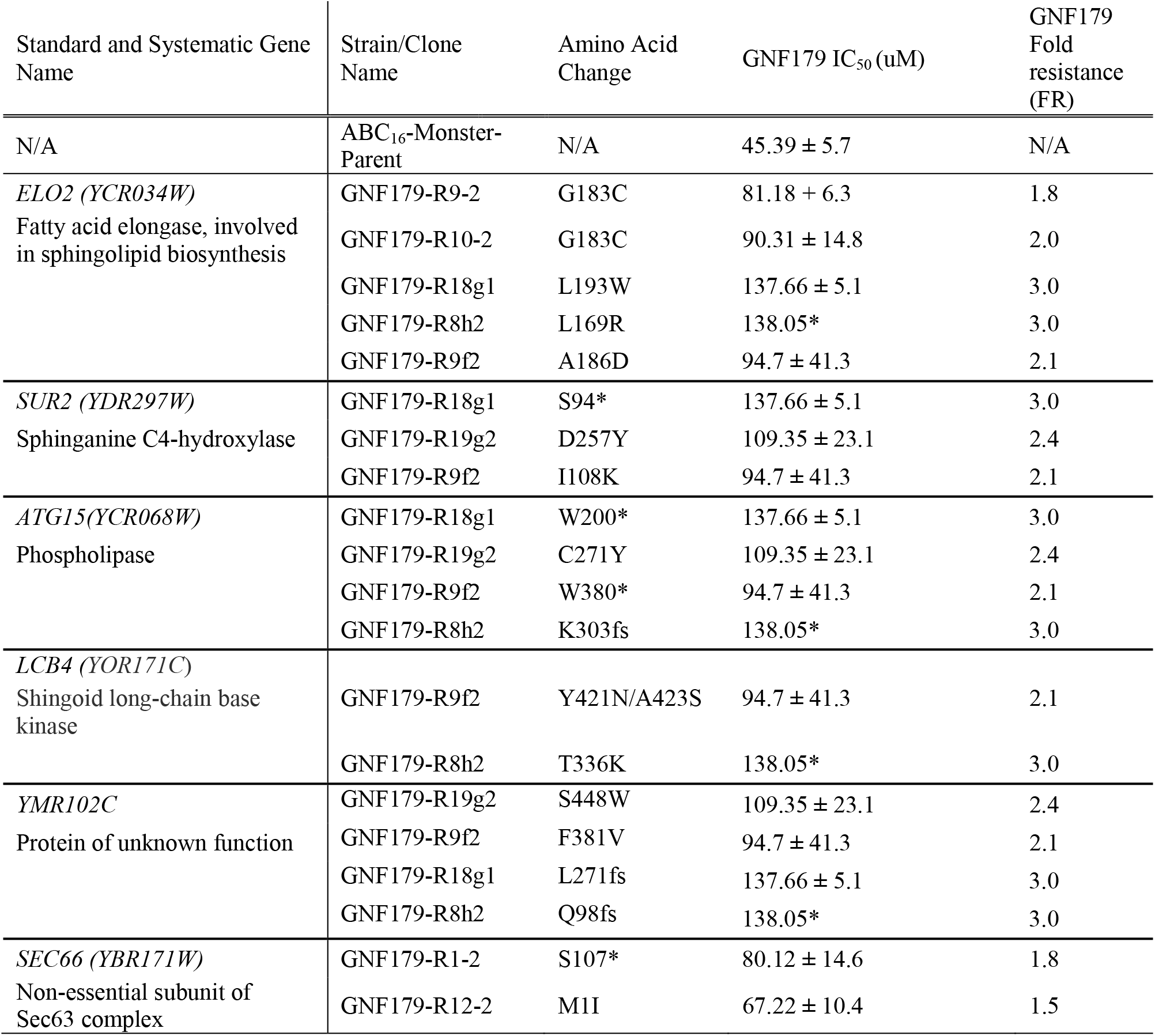
Mutations identified in more than one GNF179-resistant *S. cerevisiae* line, from a pool of 13 evolved strains. 18 hrs IC_50s_ against GNF179 (mean ± standard error with n ≥ 2) and fold resistances (calculated relative to the parental ABC_16_-Monster-Parent strain) are indicated. The complete set of coding changes for the 13 resistant lines are given in Table S1. *Replicate not available.

**Table 2.** IC_50s_ (mean and S.E.M) for GNF179 in the indicated yeast CRISPR-*Cas9* edited strains. Strain name, systematic name, 18h IC_50s_ against GNF179 (mean ± standard error with n ≥ 3) and relative change compared to unedited ABC_16_-Monster strain are indicated.

| Strain Name | Systematic Gene ID | Standard Gene Name | Mutation | Gene Description | GNF179 IC<sub>50</sub> (μM) | GNF179 Fold resistance (FR) |
| --- | --- | --- | --- | --- | --- | --- |
| ABC<sub>16</sub>-Monster | N/A | N/A | N/A | N/A | 47.30 ± 3.8 | N/A |
| EAW361 | YBR171W | *SEC66* | M1I | Non-essential subunit of Sec63 complex, involved in protein targeting and import into the ER | 70.10 ± 11.7 | 1.5 |
| EAW277 | YBR171W | *SEC66* | S107* | Non-essential subunit of Sec63 complex, involved in protein targeting and import into the ER | 74.83 ± 9.4 | 1.6 |
| EAW289 | YCR034W | *ELO2* | G183C | Fatty acid elongase, involved in sphingolipid biosynthesis; | 88.15 ± 8.7 | 1.9 |
| EAW327 | YLR372W | *ELO3* | Y307* | Fatty acid elongase, involved in sphingolipid biosynthesis; | 76.57 ± 10.0 | 1.6 |

Some of the mutations (e.g. *elo2*, *sur2*, *atg15* and *sec66*) were early stop codons and therefore resulted in truncated proteins. To provide a further layer of confirmation for the *in vitro* resistance experiments, the homozygous deletion strains (which are not attenuated like the Green Monster strain) ^31,32^ for *sec66, sur2, atg15* and *elo2* were tested against GNF179, revealing each to be 1.3-2.1-fold more resistant to IZPs (IC_50_ = 121 µM (WT) vs 188 µM (*sec66*Δ), 255 µM (*elo2Δ*) and 217 µM (*sur2Δ*) respectively). We found that the homozygous deletion strain for *yer140w* (which encodes the PfCARL homolog EMP65^33^) showed low-level resistance to IZPs (152 µM), and a homozygous deletion of *sec72*, another nonessential subunit of the complex for importing proteins into the ER, also conveyed a similar level of resistance (177 µM) (Table 3). These data suggest that modifications to either the protein export complex or lipid synthesis pathways or both can result in IZP resistance in yeast. Although the observed mutations may help overcome GNF179 treatment, it seems unlikely that these genes encode targets. Many genes are not essential and we observe stop codons. In addition, levels of resistance for isolated alleles are mild.

**Table 3.** IC_50_s (mean and S.E.M) for GNF179 in the indicated yeast haploid deletion strains. Strain name, systematic name, 18h IC_50_ against GNF179 (mean ± standard error with n ≥ 2) and relative change compared to wild-type are indicated. Strains obtained from yeast deletion collection.^31^

| Strain Name | Systematic Name | Mutation | Gene Description | GNF179 IC<sub>50</sub> (nM) | GNF179-fold resistance (FR) |
| --- | --- | --- | --- | --- | --- |
| BY4742 | N/A | N/A | N/A | 121.26 ± 2.9 | N/A |
| *sec66Δ* | YBR171W | Haploid deletion of *sec66* | Non-essential subunit of Sec63 complex, involved in protein targeting and import into the ER | 188.69 ± 11.4 | 1.6 |
| *sec72Δ* | YLR292C | Haploid deletion of *sec72* | Non-essential subunit of Sec63 complex, involved in protein targeting and import into the ER | 177.37 ± 5.1 | 1.5 |
| *elo2Δ* | YCR034W | Haploid deletion of *elo2* | Fatty acid elongase, involved in sphingolipid biosynthesis; | 254.88 ± 55.1 | 2.1 |
| *sur2Δ* | YDR297W | Haploid deletion of *sur2* | Sphinganine C4-hydroxylase; catalyses the conversion of sphinganine to phytosphingosine | 217.9 ± 12.5 | 1.8 |
| *emp65Δ* | YER140W | Haploid deletion of *emp65* | Integral membrane protein of the ER | 152.2 ± 10.5 | 1.3 |
| *atg15Δ* | YCR068W | Haploid deletion of *atg15* | Phospholipase | 137.54 ± 12.9 | 1.1 |

### *pfcarl* mutant lines show altered sphingolipid profiles

To explore sphingolipid synthesis as a direct mechanism of action, we used a HPLC (high pressure liquid chromatography) linked to triple quadrupole mass spectrometry (LC-MS/MS) to measure 19 classes of sphingolipids in *P. falciparum* Dd2 parasites exposed to 5x IC_50_ (30 nM) GNF179 for 4 hours. Of note, prior work has demonstrated mammalian-like sphingolipid biosynthetic activities in *Plasmodium* parasites^34^. Results with Dd2 WT parasites were compared to the *pfcarl* triple mutant (KAD452-R3; M81I, L830V and S1076I). LC-MS/MS analysis of saponin-lysed parasite extracts identified dihydroceramides (DH), ceramides (C), sphingosines (So), sphinganines (Sa) and sphingomyelins (SM), with varying fatty acid chains and degree of saturation (Figure 2).

**Figure 2.**
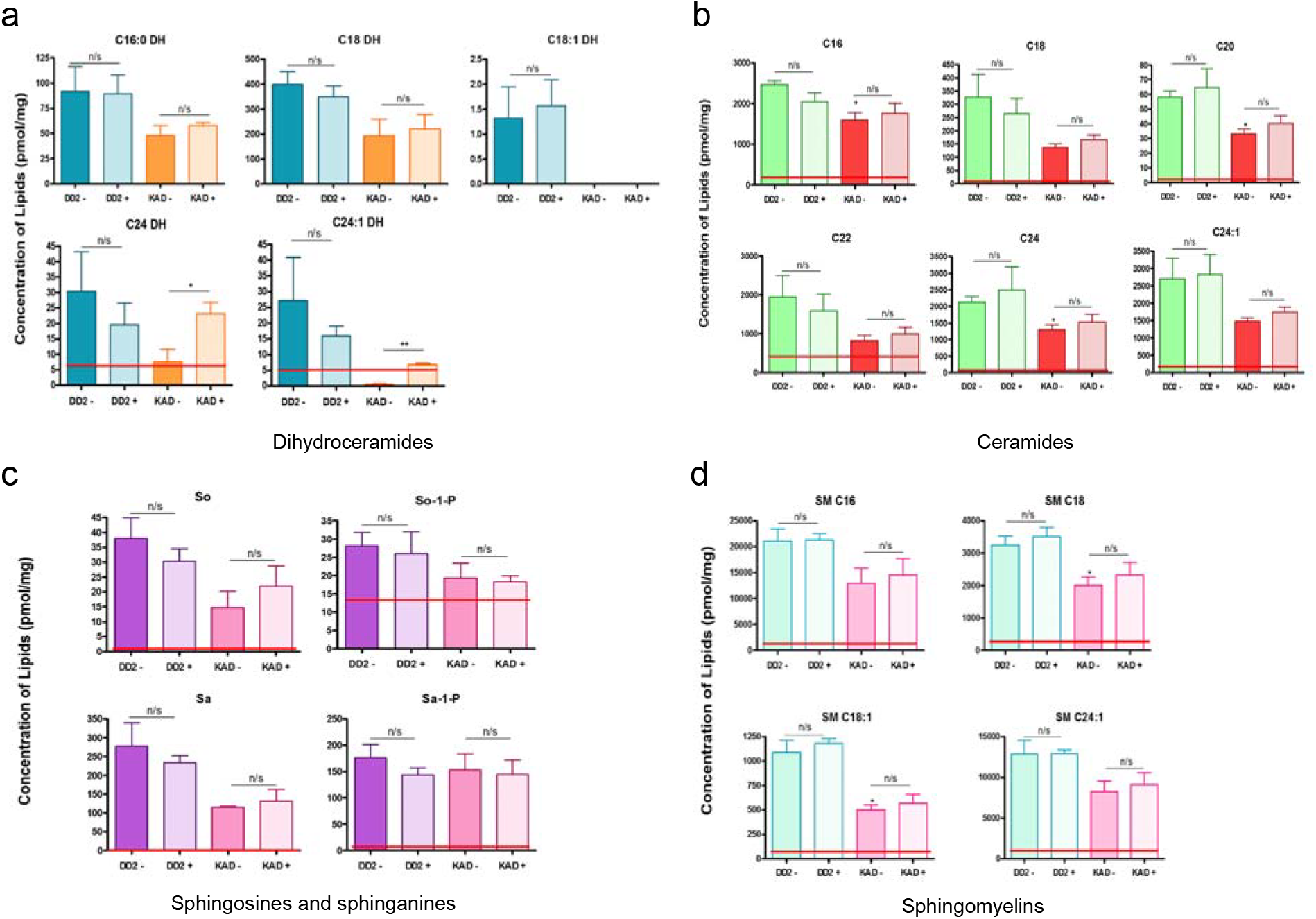
Sphingolipid profile of isolated *P. falciparum*: Wild Type and PfCARL Mutant. Composite profile of 19 sphingolipids from untreated wild type dd2 (DD2-) and *pfcarl* triple mutant (KAD452-R3: M81I, L830V and S1076I) as well as parasites treated with 25nM GNF179 (DD2+, KAD+). Lipids were normalized to protein and reported with SEM indicated by the error bars (n=3 independent experiments). Student’s t-test was employed with significance between untreated and treated wild-type and mutant indicated by solid black lines. Significance of all populations with respect to the untreated wild type is indicated by * where p < 0.05. Sphingolipid profile comprises (A) dihydroceramides (DH), (B) ceramides (C), (C) sphingosine (So), sphingosine-1-phosphate (So-1-P), sphinganine (Sa), sphinganine1-phosphate (Sa-1-P) and (D) sphingomyelins (SM) with varying fatty acid chains and degrees of saturation. Solid red line represents uninfected red blood cells as a control.

Normalized lipidomic comparison of untreated wild-type and mutant parasites indicated that the baseline sphingolipid concentration was notably lower in the *pfcarl* mutant. As an example, C18:1 dihydroceramide was undetectable in mutant parasites. Likewise, ceramide levels (Figure 2b) in untreated wild-type parasites were found to be consistently higher than in the mutant, with statistical significance being observed in 3 out of the 6 identified species (C16, C20, C24). Notably, sphingomyelins (SM C18, SM C18:1) were significantly lower compared to Dd2. These consistent differences observed in the *pfcarl* mutant line are supportive of shared mechanisms of resistance to GNF179 between *Plasmodium* and yeast.

Dihydroceramides (Figure 2a) in drug-treated WT and *pfcarl* mutant parasites tended to decrease and increase in concentration, respectively, compared to the untreated parasites. The increase in *pfcarl* mutants upon GNF179 treatment was significant for the species C24 and C24:1. Sphingosines and sphinganines (Figure 2c) demonstrated consistent trends in the sphingolipid profile in that a slight decrease was observed when wild-type parasites were treated with drug. The mutant exhibited a slightly different profile in that when treated, the sphingolipids So and Sa increased but So-1-P and Sa-1-P either remained the same or decreased slightly in concentration. None of the trends observed, however, attained statistical significance. Sphingomyelins (Figure 2d) showed no significant changes upon treatment of either WT or mutant parasites.

### Localization of GNF179 within the parasite

To gain further insight into the compound’s function we next examined subcellular localization using a fluorescently-conjugated form of GNF179 in *P. falciparum*, as recently performed for primaquine^35^. We first generated both NBD- or Coumarin-1-conjugated forms of GNF179 (Figure 3a), leveraging an existing reaction series^36^. These modified compounds retained activity against *P. falciparum* blood stages, though the potency was noticeably reduced for the Coumarin-1 conjugated form of GNF179 (19 nM for NBD and 1.2 µM Coumarin-1 vs 5 nM for non-modified GNF179). However, the GNF179-resistant strain (KAD452-R3) was also resistant to the modified forms of GNF179 (Figure 3b), indicating a similar mechanism of action for the labeled and unlabeled compounds. We first examined the localization of GNF179 in ring-stage parasites, as previous reports indicated that GNF179 is most active against early blood stages^22^. GNF179 colocalized with ER tracker Red, a live-cell dye that recognizes the ER^37^ (Figure 3c). This, combined with the *S. cerevisiae* data and the localization of the three previously identified resistance genes (*pfact*, *pfugt* and *pfcarl*) to the ER/Golgi, suggest that IZPs affect a process within this compartment.

**Figure 3.**
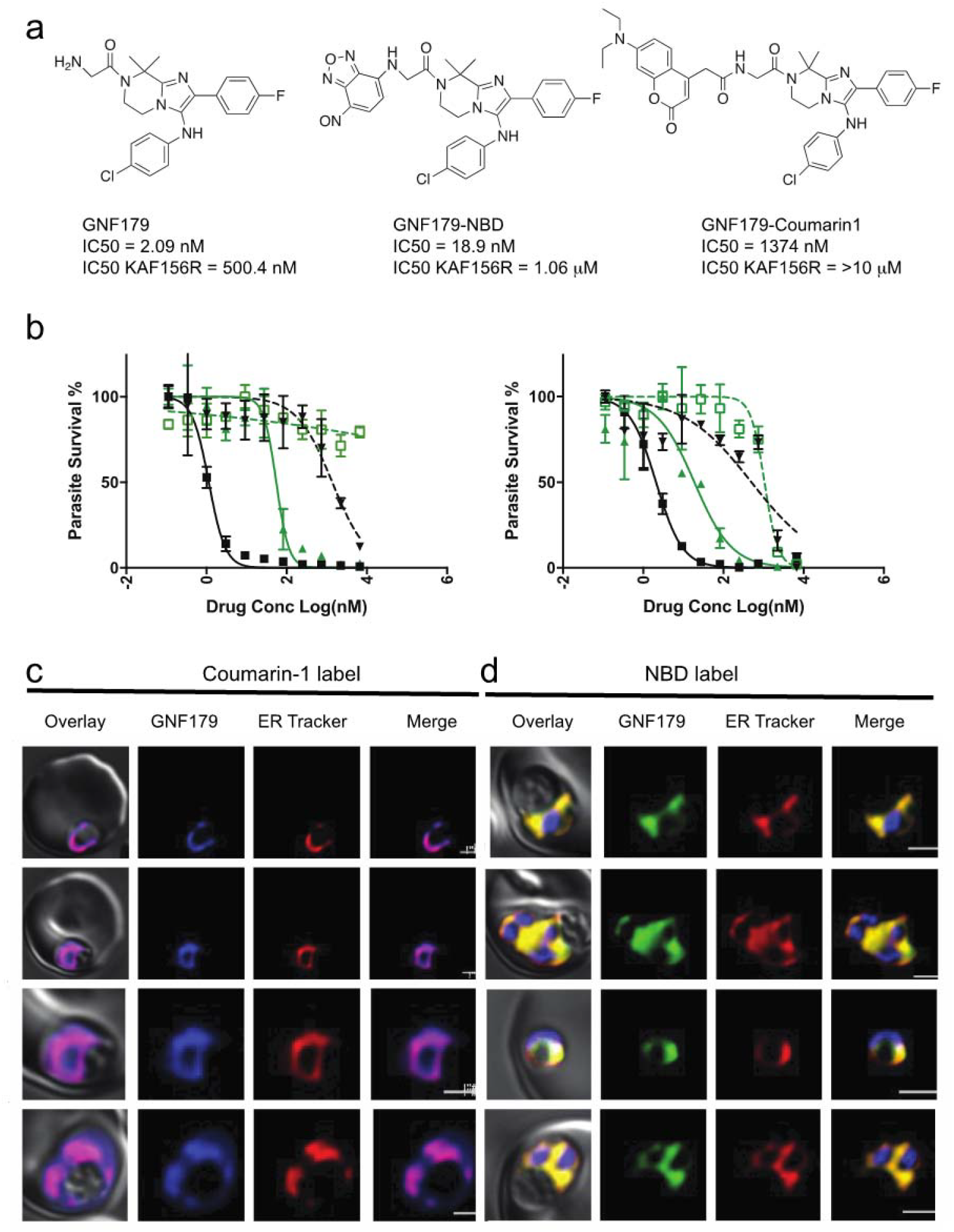
GNF179 localizes to the ER of early stage parasites and interacts with several proteins required for parasite protein trafficking into the ER. (a) Chemical structure of canonical and NBD conjugated GNF179 and Coumarin-1 conjugated GNF179. (b) Dose response curves for GNF179 and Coumarin1 (left) or NBD (right) conjugated GNF179 in wild-type and KAF156-resistant clone (KAD452-R3, containing three mutations in *pfcarl* (M81I, L830V and S1076I). (c) Colocalization of Coumarin-1 conjugated GNF179 with ER-tracker red (d) Colocalization of NBD conjugated GNF179 with ER-tracker red. Parasites are in mid-ring (6-hours post-infection) stage and were treated for 30 minutes with 2 µM GNF179-Coumarin1 and 100 nM GNF179-NBD.

### GNF179 inhibits protein export in *P. falciparum*

To further explore the hypothesis that GNF179 blocks the production or sorting of mature proteins, we next tested for synergy with known protein-export inhibitors. Brefeldin A inhibits Sec7-type GTP-exchange factors (GEFs) that catalyze the activation of a small GTPase called Arf1, responsible for ER to Golgi transport, as well as Golgi to ER retrograde transport, through the inhibition of COPI coating of secretory vesicles^38^, and is commonly used to inhibit protein secretion. We observed that treating parasites with sublethal concentrations of brefeldin A (1µM) rendered parasites 5-fold more sensitive to GNF179 (IC_50_ of 0.6 nM, vs 3 nM without brefeldin A co-treatment) (Table 4), while sensitivity to the control drug artemisinin was unchanged. This suggests that parasites exposed to GNF179 are highly sensitive to alterations in protein export.

**Table 4.**
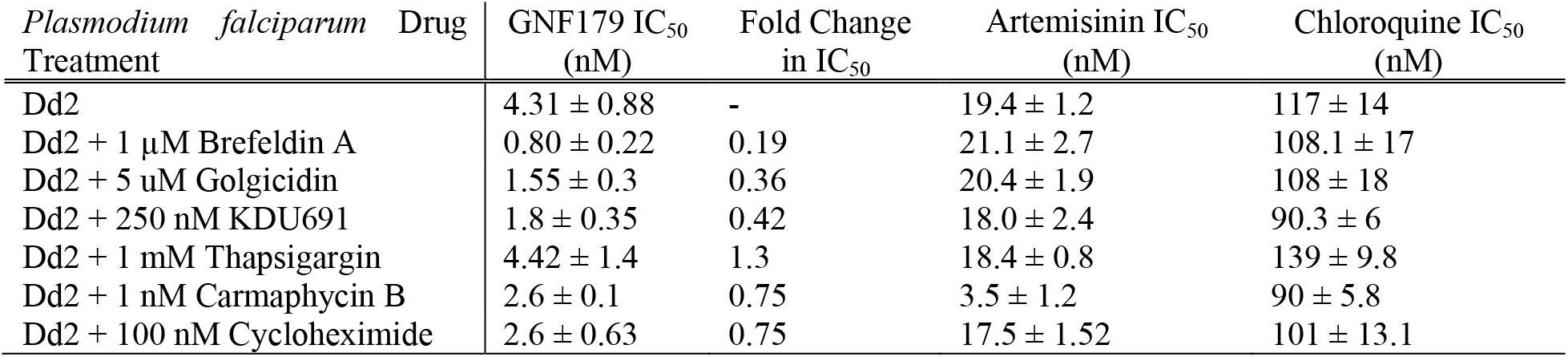
72-hour SYBR-green IC_50_s for the indicated compound combinations including GNF179. Parasites were synchronized to ring stage before IC_50_s were measured using the SYBR green method. IC_50_s presented as Mean ± Standard Error with ≥2 biological replicates of two technical replicates each.

**Table 5.**
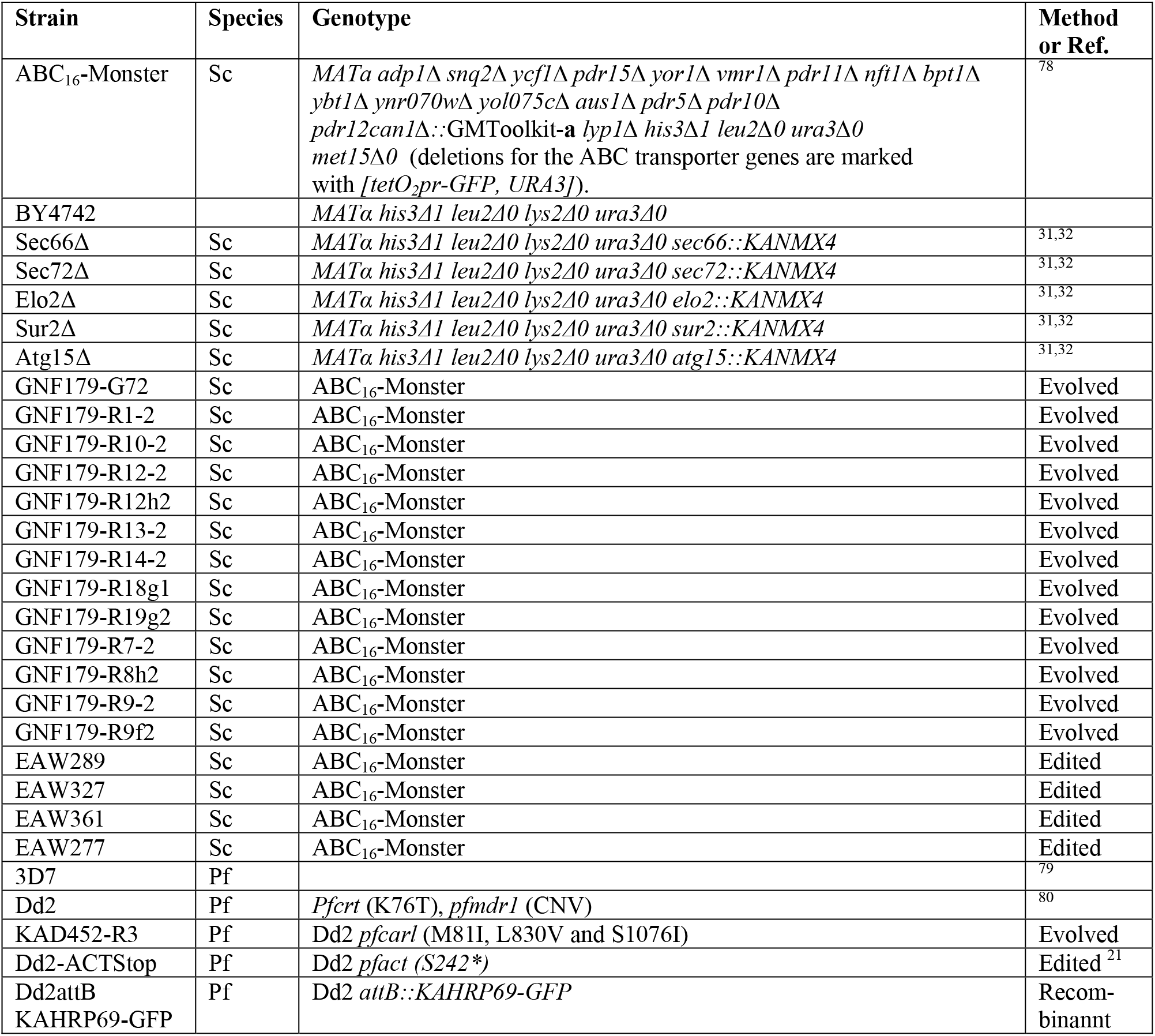
Strains used in this study. Sc, *S. cerevisiae*, Pf, *P. falciparum*.

A similar effect was observed for golgicidin, another inhibitor of protein export that works through inhibition of Golgi function^39^. Simultaneous treatment with 5 µM golgicidin (IC_50_ 11µM) also rendered parasites 2-3-fold more sensitive to GNF179 (Table 4). Importantly, this effect was specific to GNF179, rather than merely a sign of generalized parasite growth impairment, as artemisinin, atovaquone and chloroquine all showed no synergy with any of these three compounds. This synergistic effect was exclusive to inhibitors of protein secretion from the ER and Golgi, as simultaneous treatment with thapsigargin, which inhibits PfATP6 and causes depletion of ER calcium levels leading to downstream inhibition of ER chaperones^40^, caused no change in GNF179 potency. In addition, we tested for synergy between GNF179 and inhibitors of the *P. falciparum* proteasome (Table 4), to determine whether the identified increase in ubiquitinated proteins was directly involved with the efficacy of GNF179 or rather was a downstream consequence. We did not observe any synergy between GNF179 and carmaphycin B, a natural product-derived proteasome inhibitor^27^. This is in contrast to artemisinin, which increases the potency of carmaphycin B. This indicated to us that the increase in protein ubiquitination was a secondary effect of GNF179 inhibiting protein trafficking, rather than a direct cause.

To examine whether protein secretion was being blocked, we sequentially looked at protein processing in *P. falciparum*. In free-living organisms, proteins are trafficked to membranes or to the extracellular environment. In intracellular malaria parasites, however, proteins can be further exported into the infected red blood cell (RBC) or hepatocyte via the PTEX complex, a parasite-specific secretory complex located in the parasitophorous vacuole membrane^41^. To examine these two processes, we used two different transiently-expressed reporters. The first reporter (Figure 4a) is based on a PfEMP3-GFP fusion protein and includes the first 82 amino acids of PfEMP3 (PF3D7_0201900) fused to GFP. *pfemp3* encodes *P. falciparum* Erythrocyte Membrane 3, a protein that is exported to the surface of the infected RBC and that is needed to form knobs that permit cellular adhesion to vascular endothelial cell surface receptors. These 82 amino acids of PfEMP3 include both the signal peptide and PEXEL motif, a 5 amino acid sequence (RxLxE/Q/D) present in most *Plasmodium* proteins exported to the RBC cytosol. This sequence is cleaved by Plasmepsin V, a type I integral membrane-bound protease with the active domain located on the luminal side of the endoplasmic reticulum (ER).^42^ Based on where the export pathway is blocked three different potential protein products may be observed, the full length fusion protein, the PEXEL-domain cleaved protein, and mature GFP. As a control, we used WEHI-842, a peptide mimetic Plasmepsin V inhibitor (EC_50_ = 400 nM)^42^. Treatment with WEHI-842 showed a marked accumulation of the full-length, unprocessed protein relative to the untreated control, as expected, given the role of Plasmepsin V in cleavage of PEXEL signal sequences. Parasites treated with high doses of GNF179, on the other hand, showed a marked decrease in levels of all forms of the secreted reporter construct, with little evidence of cleavage (Figure 4a). In contrast, levels of HSP70 protein, which does not have a signal sequence, were unchanged after treatment with GNF179 or WEHI-842. This finding suggests that proteins trafficked through the ER, but not cytoplasmic proteins, are being degraded or potentially not being synthesized due to exposure to GNF179.

**Figure 4.**
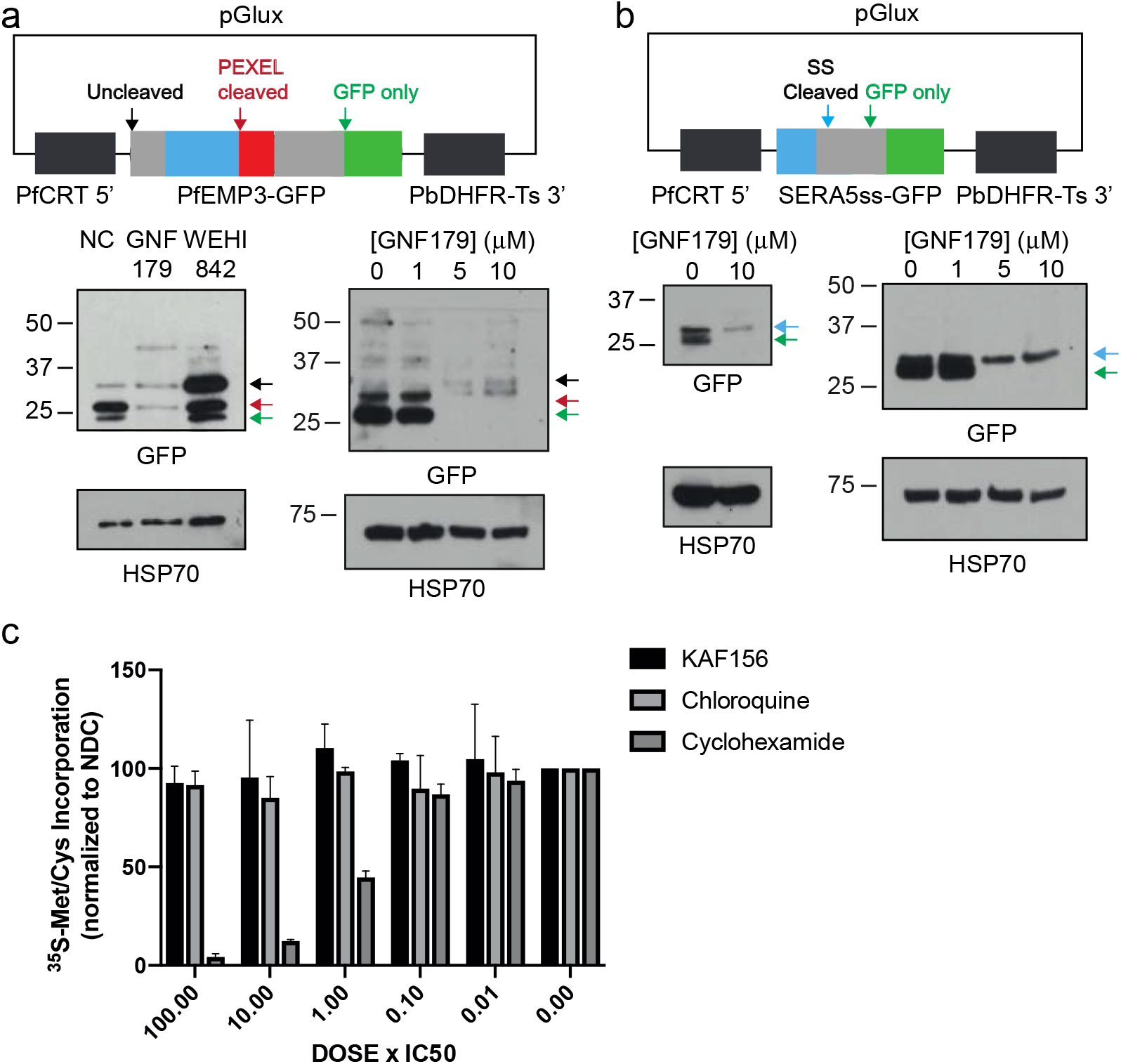
Secretion reporter constructs demonstrate that GNF179 inhibits protein export of *Plasmodium* falciparum. (a) Protein expression levels of PfEMP3-GFP reporter. This fusion includes the signal peptide and PEXEL motif of PfEMP3. By immunoblot, 3 protein products are seen with anti-GFP antibodies. The three indicated bands, in response to probing with GFP, as follows: 1. Full length protein (Black arrow), 2. PEXEL cleaved protein (red arrow), 3. GFP degradation product (green arrow). HSP70 is used as a loading control. (b) SERA5ss-GFP fusion reporter treated with GNF179. By immunoblot with anti-GFP antibodies we see two protein products for this construct: 1. Signal peptide cleaved (blue arrow) and 2. GFP degradation product (green arrow). HSP70 serves as a loading control. (c) ^35^S incorporation of newly synthesized amino acids at different concentrations of KAF156, chloroquine (negative control for inhibition) and cycloheximide (positive control for inhibition). Counts were normalized to data obtained from no-drug controls (NDC).

The second reporter construct was a SERA5ss-GFP fusion, containing the signal peptide of SERA5 (PF3D7_0207600) fused to GFP (Figure 4b). SERA5 is an exported serine protease that plays a role in parasite egress from the infected RBC^43^. This fusion protein is co-translationally inserted to the ER and the signal peptide is removed by a signal peptidase, leading to trafficking of the mature protein to the parasitophorous vacuole (PV)^43^. SERA5ss-GFP is therefore used as a marker of protein secretion to the PV, versus the infected RBC. Without treatment, we observed two protein products for this chimera using GFP antibodies. After exposure to GNF179, the reporter showed a marked decrease in overall abundance relative to the HSP70 control, as observed for PfEMP3 reporter above (Figure 4b). In addition to marked overall reduced expression, we also observed an accumulation of the unprocessed reporter protein, based on the overall ratio of the two products. Based on results with these two reporter systems, we speculated that there was an impairment of protein secretion that occurs before signal peptide cleavage, leading to degradation of the uncleaved products or potentially a block in synthesis. Although this could be a result of reduced total protein synthesis, we observed continued incorporation of ^35^S methionine after treatment with GNF179, in contrast with cycloheximide. Thus, if protein synthesis was inhibited, it was likely to only be for a subset of proteins. To test whether the loss of reporter signal via western blot could be due enhanced proteolysis through the ubiquitin system we compared the effects of MG132, a proteasome inhibitor, on cleaved and uncleaved reporter levels both in the presence and absence of GNF179. These data showed that proteasome inhibition, in contrast to GNF179 inhibition, did not affect the levels or processing of either the PfEMP3 or the SERA5 reporter and that there was no observed synergy with GNF179 (Figure S1). Interestingly, increases in total ubiquitinated proteins were observed after both MG132 and GNF179 treatment, although banding patterns were somewhat different suggesting that different classes of proteins could be ubiquitinated after treatment with the two compounds.

We also tested whether native parasite proteins that are normally exported to the surrounding infected RBC would be retained within the parasite after exposure to GNF179. We therefore examined the intracellular levels of three PEXEL-containing proteins: PTP2 (PF3D7_0731100), PIESP2 (PF3D7_0501200) and SERA5 (PF3D7_0207600). In all three cases, after a short, 3-hour exposure with 10µM GNF179 (same conditions as above for the reporters for both GNF179 and WEHI-842) (Figure S2), we observed a modest accumulation of all three proteins suggesting impairment of protein export from the parasite (Figure S2). A conditional knockdown of PfSEC62 using the glmS ribozyme system^44,45^ showed that parasites were 3-fold more sensitive to GNF179 (0.66 nM for 3D7 wildtype vs 0.24 nM for PfSEC62 kd) (Figure S3).

Finally, we validated the effect of GNF179 on parasite protein export in live parasites through two parallel strategies, using brefeldin A (BFA) as a positive control: visualization of export of fluorescent reporter proteins, and examination of functional phenotypes that would result from inhibition of protein secretion. To begin, we constructed a parasite strain that bears a fusion between the Knob-Associated Histidine Rich Protein and GFP^44^ (Figure 5a). The chimeric gene, which bears the first 69 amino acids of KAHRP containing the signal peptide (SP) and PEXEL motif (Px), was expressed from a *pfcrt* promoter and was integrated into the *cg6* locus in Dd2-attB parasites using the attP × attB integrase system^45^. In the absence of GNF179 the GFP reporter was trafficked to the parasitophorous vacuole (PV) as well as to the RBC cytosol, as shown by the GFP staining in the PV surrounding the parasite (Figure 5b). In contrast, treatment with brefeldin A resulted in accumulation of the GFP reporter in the parasite ER as evidenced when co-staining with an anti-PDI antibody (cyan) or by using ER tracker red (Figure 5b and Figure S4). The white dotted outlines indicate overlap of green and cyan labels. To better understand in which subcellular compartment the export block occurred we co-stained with antibodies to both the cis-Golgi resident protein, ERD2,^46^ and to the ER (PDI)). In contrast to BFA - treated parasites, images of parasites treated with 5x IC_50_ GNF179 (25 nM) for 16 hours showed colocalization of the GFP signal at times with both organelles (Figure 5c), although we found more colocalization with the ER than with the Golgi (area circles with white dotted lines). Interestingly, staining of parasites with ER-tracker after GNF179 treatment showed an expansion of ER tracker positive space relative to nuclear staining (Figure S4 a-c). 3D volume quantification of the ratio between DAPI-positive staining to ER-staining in live, GNF179-treated parasites showed a statistically-significant expansion of the ER (Figure S4d).

**Figure 5.**
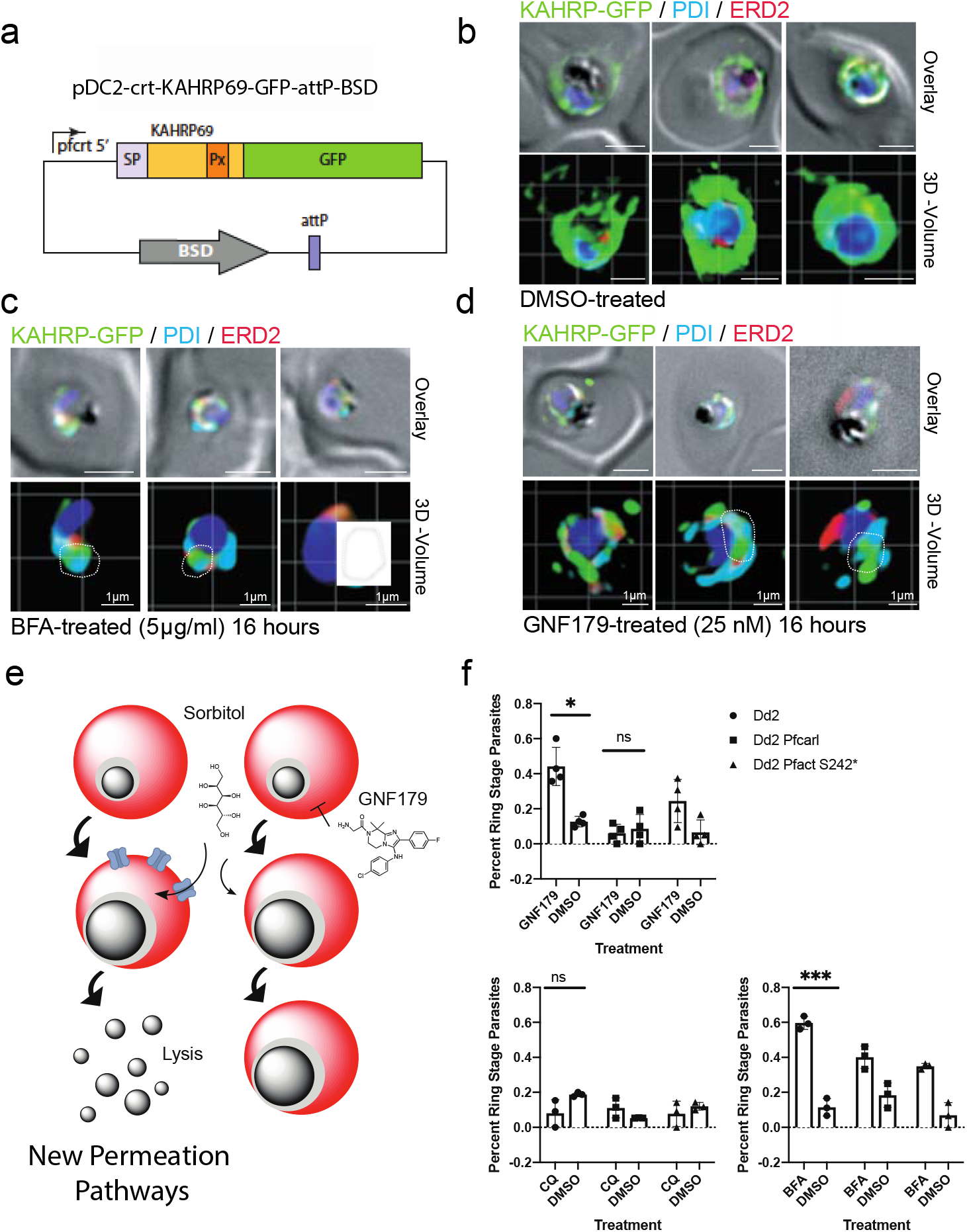
Confirmation of GNF179’s inhibition of protein export in *P. falciparum* blood stage parasites. (a) Vector used for assays shown in b, c., d. (b, d) Imaging of GFP reporter to the plasma membrane under the indicated compound treatments for 16 hours. Parasites were fixed and stained with Hoechst 33342 (blue), α-GFP (green), α-ERD2 (red) and α-PDI (cyan) antibodies. Scale bars: 2 µm unless otherwise indicated. Overlay: DIC images merged with fluorescent channels. (e) Model for establishing new permeation pathways. If proteins are not exported to the red cell where they can form new permeation pathways, sorbitol will not be imported and cause lysis. (f). Percentage of trophozoites from an asynchronous parasite population 24 hours after sorbitol synchronization, treated as indicated by GNF179 (5nM), Chloroquine (CQ, 500nM) or brefeldin A (BFA, 5µM). Statistical significance was determine using a paired, two-tailed t-test. Dd2 *pfcarl* experiments were conducted with the *pfcarl* evolved triple mutant (KAD452-R3)) and edited Dd2-ACTStop mutant. For sensitive parasites (Dd2), 25nM of GNF179 was used, while 1µM was used on resistant parasites.

To functionally confirm a secretion defect, we next determined whether parasites treated with GNF179 would be less sensitive to sorbitol treatment, which kills mature parasites and is used to synchronize cultures as only ring-stage parasites survive. For sorbitol to kill later stage parasites, the new permeability pathways (NPPs) must first be established in the RBC, which requires the export of a network of transport proteins ^49^ (Figure 5e). Here, we observed that 4 hours of pretreatment with 25 nM GNF179 led to a loss of sensitivity to sorbitol, as seen via the presence of a nearly asynchronous culture 24 hours post synchronization (Figure 5f). This was in contrast to parasites that were GNF179 resistant, resulting from mutations in either *pfcarl* (KAD452-R3, Figure 5f) or, to a lesser degree, *pfact* (Dd2-ACTStop, Figure 5f). Those resistant parasites showed less protection and more sorbitol lysis. As controls we observed a similar phenotype with brefeldin A but not with chloroquine. The lack of sorbitol sensitivity in the presence of GNF179 further highlights the impairment of ER-mediated protein trafficking due to GNF179 exposure.

## Discussion

Once the parasite has successfully invaded the host cell, the parasite undergoes a rapid induction of protein synthesis^50^. This includes the folding and sorting of hundreds of membrane proteins and the secretion of hundreds of proteins into the host cell via the parasite ER and the PTEX complex in the PVM, leading to a dramatic rearrangement of host cell processes, all essential for parasite survival. Proteins needed for these essential processes may make effective drug targets, as shown by the development of WEHI-842, an inhibitor of Plasmepsin V^51^. In addition, KDU691 targets PI4K^52^, a protein involved in a variety of processes including vesicular trafficking from the Golgi apparatus. While the trafficking of proteins into the infected RBC may only occur during the asexual blood stage, general trafficking and protein modification is a process that occurs, at least to some degree, at all stages of parasite development. Therefore, proteins that play a role in protein processing or sorting pathways would be logical targets for the type of multistage (hepatic, asexual blood and gametocyte stages) activity possessed by KAF156.

The ER serves as the initial entry into the protein trafficking and secretion pathways, as a site for protein folding, and as the location of a significant portion of the post-translation modification of parasite proteins. All of our available data suggest that KAF156 exerts its effect in this organelle, including the localization of gene products that give resistance (in both *S. cerevisiae* and *P. falciparum*) to KAF156/GNF179 as well as the localization of NBD and Coumarin-labeled compounds. The ER is responsible for many functions, including lysine acetylation, protein folding, glycosylation and sphingolipid biosynthesis (Figure S5). After proteins have been modified and folded, they are sorted to other compartments. While KAF156 may block the sorting of proteins to other compartment, our data seem more compatible with this agent working at an earlier stage, protein-production stage, possibly during protein folding: Western blot analysis shows a substantial reduction in the total amounts of secreted reporter proteins relative to cytoplasmic proteins after GNF179 treatment, suggesting that the synthesis of both membrane and PTEX-trafficked proteins may be disrupted. The reduction in protein levels is in contrast to using the WEHI-842 inhibitor of the ER-based Plasmepsin V export protease, where a shift from the processed to the unprocessed form of the protein is observed yet protein reporter levels stay high. Levels of proteins that are not exported to the RBC but have a simple signal sequence are also reduced (Figure 4).

While a block in the synthesis of membrane proteins might be expected to result in an unfolded protein response, and synergy with proteasome inhibitors such as carmaphycin B, there are different degradation pathways for cytoplasmic proteins versus proteins that are destined to be secreted. Recent data reveal the existence of two pathways for dealing with protein aggregates in the ER: ERAD(I) and ERAD(II)^53^. The second pathway depends on autophagy and lysosomal trafficking for the disposal of large protein aggregates, and is observed in human cells when Acetyl-CoA Transporter is disrupted^54^. It is likely that ERAD(II) is used when secreted reporters (e.g. KAHRP69-GFP) are overexpressed, as we observed no great increase in ubiquitinated proteins after GNF179 treatment except when a proteasome inhibitor (MG132) was used in combination with GNF179 (Figure S1). An expansion of the ER, similar to that observed after GNF179 treatment in malaria parasites (Figure S4), is observed when yeast cells are treated with agents that induce the unfolded protein response. ER expansion induces autophagy as a mechanism to deal with ER-stress and eliminate the blockage^55^.

Although GNF179 appears to work in the ER, the exact target of GNF179 is not known. On the other hand, the cancer and antinfectives drug discovery communities have explored this organelle and its processes for targets (reviewed in ^56^). The ER protein-folding factory examined here also holds targets for antivirals as viruses need to be packaged within the ER^56^. In addition, it has been established that the *P. falciparum* Signal Peptide Peptidase PfSPP1 is a drug-able target and its inhibition results in parasite death across the lifecycle^57^. Conservation of critical targets across species seems to be more the rule than the exception although selectivity of inhibitors may be very different.

Although it is difficult to predict a target based on chemical structure, KAF156 has structural features that suggest it may inhibit a kinase or other ATP or GTP binding proteins. Many potential essential targets with ATP or GTP binding pockets are present ER-dependent protein packaging pathway. Members of the Sec62 translocon complex could be possible targets. However, Sec62 does not have an obvious catalytic site, a feature that is associated with many targets, although it is possible that IZPs are allosteric inhibitors. Signal recognition particle (SRP) is a protein complex critical for the translocation of proteins into the ER ^*58*^. Within this complex, both SRP54 and SRPα harbor GTPase domains, both of which would represent a prime drug-able target. However, these data would not explain the increased ER volume that is observed with GNF179-exposed parasites. Other candidates may include heatshock proteins such as the GRP94-like protein endoplasmin (PF3D7_1222300). This essential protein has an ATP-binding domain and selective targeting with small molecule inhibitors is possible^59^. GRP94 homologs are not found in yeast and bacteria. However, GRP94 is somewhat homologous to HSP82. In fact, in our yeast selections we detect a Ser to Ile substitution at position 17 that lies in proximity to the ATP-binding pocket. However, the lack of a close ortholog may explain the 10,000-fold difference in potency in the yeast model. In addition, binding immunoglobulin protein (BiP), the Hsp70 homologue within the ER (PF3D7_0917900), requires ATPase activity to perform the final step of protein translocation from the SEC61 translocon into the ER^60^ and could also be a plausible target. Chemical targeting of BiP is feasible.^61^ The hypothesis that KAF156/GNF179 targets protein folding in the ER is supported by the role of EMP65 as a guardian factor that protects newly synthesized, unfolded polypeptides from degradation (See Figure S5 for model). If *pfcarl-emp65* is mutated, more premature degradation may occur in the presence of GNF179/KAF156, which may allow the cells to tolerate 10X more GNF179.

Another potential class of proteins are Rab GTPases. Given the synergy we observed between GNF179 and brefeldin A, GNF179 could inhibit COPI-mediated transport from the ER to the Golgi, similar to the mechanism of action of brefeldin A^62^. Sar1, an essential gene involved in the Sec23 complex, for example, is a small GTPase that functions as a target in a yeast brefeldin suppressor screen^62^. The lysine transferase that adds acetyl groups to nascent peptides, helping them fold and which relies on PfACT activity could be a target.

Protein glycosylation in the ER may could also be inhibited and may lead to unfolded aggregates within the ER. Tunicamycin, a nucleoside analog, inhibits protein N-linked glycosylation by preventing core oligosaccharide addition to nascent polypeptides by binding to UDP-N-acetylglucosamine--dolichyl-phosphate N-acetylglucosamine phosphotransferase (Alg7). Tunicamycin thereby also blocks protein folding and transit through the ER. The *P. falciparum* version of Alg7, PF3D7_0321200, could theoretically be a target, although the structure of KAF156 is not similar to tunicamycin.

It is although worthwhile noting that several additional compounds may work via similar or related mechanisms. For example, mutations in *pfcarl* are acquired when parasites are treated with sublethal concentrations of MMV007564^63^. In particular, if there is a *pfcarl* or *pfact* resistance mechanism, it seems likely that protein folding or trafficking in the ER is targeted. More work will be needed to determine whether MMV007564 has the same target as KAF156 or if they are only in the same pathway. By comparison, there are a variety of different chemotypes that inhibit mitochondrial function^64^, as well as different chemotypes that interact with the same active site, however there still seem to be a limited number of high-value inhibitor binding sites in this pathway, and they appear repeatedly.

An additional mechanism of action for IZPs could be inhibiting the process of ER autophagy. The mammalian orthologue of Sec62, potentially a target of GNF179, is required for the degradation of excess ER components in a process termed ER autophagy^65^. *In vitro* directed evolution of *S. cerevisiae* using GNF179 identified mutations in the *atg15* and *atg22*, genes that play a role in ER-autophagy. In addition, GNF179 induced an expansion in ER size upon treatment of *P. falciparum*. Collectively, these results suggest that IZPs affect ER homeostasis and function by inhibiting proteins critical for ER-phagy.

Strategies for finding the target of KAF156 remain limited in the absence of genetic methods, and few antimalarials have been matched with their target when genetic methods fail. Affinity-based methods were used to match MMV030048 to PI4K with success^66^, although such strategies required the use of a modified ligand. Cellular Thermal Shift Assays (CETSA) methods that involve incubating total cellular extracts with an unmodified ligand and identifying those proteins that resist heat denaturation in the presence of compound is a strategy that shows promise^67^. Another potential option involves a genome-wide knockdown or knock-in libraries, which have been used with success in trypanosomes and to find the target of cladosporin also using the *S. cerevisiae* model^68^. A limitation of all of these genome-wide methods is that they may produce hundreds of possible candidate molecules, and it can be challenging to sort through them using the slow methods available in *Plasmodium*. While finding the exact target may require some work, the studies here should allow prioritization of candidates that are identified in genome-wide experiments.

## Materials and Methods

### *In vitro* Resistance Evolution and Whole Genome Sequencing of GNF179-resistant *S. cerevisiae*

Sublethal concentrations of GNF179 were added to 50ml conical tubes containing 20µl of saturated *S. cerevisiae* ABC_16_-Monster cells in 20ml of YPD media. Each selection was cultured under vigorous shaking until the culture reached saturation. Saturated cultures were diluted into fresh YPD media containing increasing GNF179 concentrations, and multiple rounds of selection were performed. Cells of cultures that were able to grow in substantially higher drug concentrations than the parental cell line, were streaked onto agar plates containing GNF179 to select for single colonies. Single colonies were isolated, and IC_50_ assays, prepared by two-fold dilution were performed to determine the degree of evolved resistance vs. that of the parental strain.

Genomic DNA (gDNA) was extracted from yeast samples using the YeaStar Genomic DNA kit (Cat. No D2002, ZYMO Research). Sequencing libraries were prepared using the Illumina Nextera XT kit (Cat. No FC-131-1024, Illumina) following the standard dual indexing protocol, and were then sequenced on the Illumina HiSeq 2500 in RapidRun mode to generate paired-end reads 100bp in length. Reads were aligned to the *S. cerevisiae* 288C reference genome (assembly R64) using BWA-mem^69^ and further processed using Picard Tools (http://broadinstitute.github.io/picard/). A total of 13 clones were sequenced to an average coverage of 47.3x, with an average of 99.3% of reads mapping to the reference genome. SNVs and INDELs were called using GATK HaplotypeCaller, filtered based on GATK recommendations^70^ and annotated with SnpEff^71^. Variants were further filtered by removing mutations that were present in the both drug-sensitive parent strain and resistant strains, such that mutations were only retained if they arose during the drug selection process.

### CRISPR-*Cas*9 Allelic Exchange in *S. cerevisiae*

CRISPR-*Cas*9 genome engineering was performed on the *S. cerevisiae* ABC_16_-Monster strain using vectors p414 and p426 obtained from the Church lab (Addgene) as previously described^72^. To produce gRNA plasmids specific to the desired mutation sites, oligonucleotides were synthesized (Integrated DNA Technologies) to match the target sequence and contain a 24 base-pair overlap with the p426 vector backbone. Gene-specific gRNAs were amplified by PCR, transformed into Stellar^TM^ competent *E. coli* cells (Takara) and selected on LB-Ampicillin plates. DNA was isolated from transformed *E. coli* cells and purified using the QiaQuick Miniprep kit (Qiagen) and quantified via Qubit Fluorometric Quantitation (ThermoFisher). Cas9-expressing ABC_16_-Monster cells were transformed with 300-500ng of gene-specific gRNA vector and 1-2 nmole of synthesized donor template (IDT) containing the desired base-pair substitution via standard lithium acetate method. Transformed cells were selected on methionine and leucine deficient CM-glucose plates. Each mutation was confirmed with Sanger sequencing (Eton Bioscience).

### *P. falciparum* culture

*P. falciparum* Dd2 strain parasites were cultured under standard conditions^73^, using RPMI media (Thermo Fisher # 21870076) supplemented with 0.05 mg/ml gentamycin (Thermo Fisher # 15710072), freshly-prepared 0.014 mg/ml hypoxanthine (Sigma Aldrich #H9377), 38.4 mM HEPES (Sigma Aldrich #H3375), 0.2% Sodium Bicarbonate (Sigma Aldrich #S5761), 3.4 mM Sodium Hydroxide (Sigma Aldrich #S8045), 0.05% O+ Human Serum (Denatured at 56°C for 40 min and obtained from Interstate Blood Bank, Memphis, TN) and 0.0025% Albumax (Thermo Fisher Scientific # 11021037). Human O+ whole blood was obtained from The Scripps Research Institute (La Jolla, CA) using the Normal Blood Program and Humann Subjects Protocol Number (IRB-12-5933). Leukocyte-free erythrocytes were stored at 50% hematocrit in RPMI-1640 screening media (as above, but without O+ human serum and with 2x albumax concentration) at 4°C for one to three weeks before experimental use. Cultures were monitored every one to two days via direct observation of parasite infection using light microscopy-based observation of Giemsa-stained thin blood smears of parasite cultures. Specific parasite cultures used are as indicated in the specific experiments.

### Conjugation of Coumarin-1 and NBD with GNF179

GNF179 was conjugated with a coumarin-1 fluorophore as previously described^36^. Briefly, Meldrum’s acid was acylated with methyl 5-chloro-5-oxovalerate and subsequently treated with methanol to provide coumarin β-keto ester. Next, the β-keto ester was first reacted with resorcinol under acidic conditions and then hydrolyzed with lithium hydroxide to provide 4-(7-hydroxy-2-oxo-2H-chromen-4-yl) coumarin butanoic acid 2.8 Finally, GNF179 and coumarin butanoic acid was coupled under standard EDCI/DMAP coupling conditions to yield the probe Coumarin-1-GNF179. To construct the NBD modified version, GNF179 was conjugated with a nitrobenzoxadiazole (NBD) fluorescent label by reacting GNF179, triethylamine, and commercially available NBD-Cl in dimethylformamide.

All reactions were performed in flame- or oven-dried glassware sealed with rubber septa and under nitrogen atmosphere, unless otherwise indicated. Air- and/or moisture-sensitive liquids or solutions were transferred by cannula or syringe. Organic solutions were concentrated by rotary evaporator at 30 millibar with the water bath heated to not more than 50°C, unless specified otherwise. Thin-layer chromatography (TLC) was performed using 0.2 mm commercial silica gel plates (silica gel 60, F254, EMD Chemicals). Nuclear Magnetic Resonance (NMR) spectra were recorded on a Varian (^1^H NMR: CDCl_3_ (7.26) at 600 MHz; ^13^C NMR: CDCl_3_ (77.16) at 151 MHz). All spectra were taken in CDCl_3_ with shifts reported in parts per million (ppm) referenced to protium or carbon of the solvent (7.26 or 77.16, respectively). Coupling constants are reported in Hertz (Hz). Data for ^1^H-NMR are reported as follows: chemical shift (ppm, reference to protium; s = single, d = doublet, t = triplet, q = quartet, dd = doublet of doublets, m = multiplet, coupling constant (Hz), and integration). High Resolution Mass Spectra (HRMS) were acquired on an Agilent 6230 High Resolution time-of-flight mass spectrometer and reported as m/z for the molecular ion [M+H]+.

### GNF179-1-Coumarin

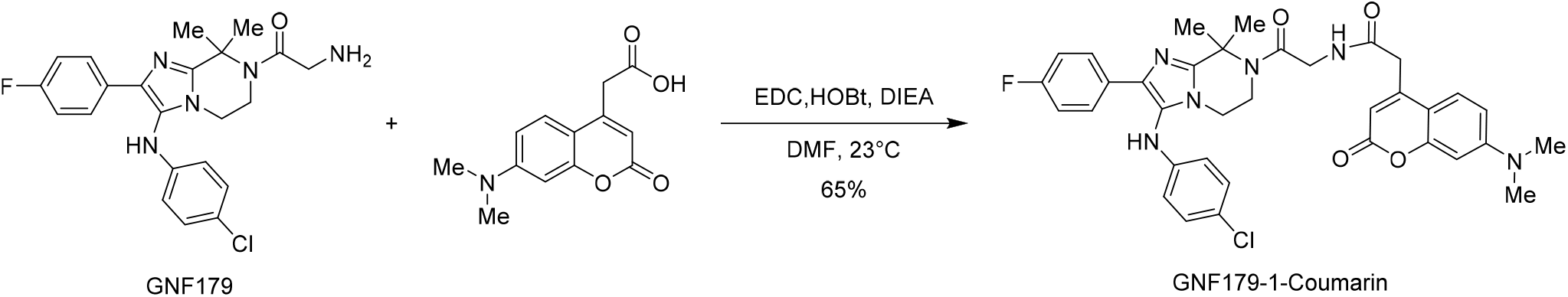

#### GNF179-1-Coumarin

A dry 5 mL round-bottom flask equipped with a stir bar, sealed with a rubber septum was charged with 7-dimethylaminocoumarin-4-acetic acid (4 mg, 0.016 mmol, 1.0 equiv.) and GNF179 (7 mg, 0.016 mmol, 1.0 equiv.) in dry DMF (0.2 mL). The solution was cooled to 5 °C and solid EDC (4 mg, 0.023 mmol, 1.42 equiv.), HOBt (3 mg, 0.019 mmol, 1.2 equiv.) and neat DIEA (3 μL, 0.019 mmol, 1.2 equiv.) were added to the mixture. The solution as stirred for 30 mins and the reaction was allowed to warm to ambient temperature and react for 18 hours. The reaction was then diluted with EtOAc (3 mL) and washed with water (1 mL). The mixture was extracted with EtOAc (2 mL). The combined organic layers were dried over sodium sulfate, filtered, and concentrated. The crude product was purified by silica column chromatography, eluting with Hexanes: EtOAc (10: 90) affording the title compound (6 mg, 9.13 mmol, 63%) as yellow color solid.

#### NMR results

**R**_**f**_ = 0.5 (silica gel, 100 EtOAc); ^**1**^H NMR (600 MHz, CDCl_3_) δ 7.73 (dd, *J* = 8.5, 5.6 Hz, 2H), 7.40 (d, *J* = 9.0 Hz, 1H), 7.13 (d, *J* = 8.6 Hz, 2H), 6.95 (m, 3H), 6.56 (m, 3H), 6.35 (d, *J* = 2.1 Hz, 1H), 6.01 (s, 1H), 5.79 (s, 1H), 4.06 (d, *J* = 3.7 Hz, 2H), 3.68 – 3.64 (m, 2H), 3.63 (s, 2H), 3.47 (m, 2H), 3.00 (s, 6H), 1.88 (s, 6H). ^**13**^C NMR (151 MHz, CDCl_3_) δ 167.98, 167.27, 162.70, 162.02, 161.07, 155.94, 153.11, 149.83, 147.44, 144.32, 133.72, 129.74, 127.64, 127.59, 125.60, 124.46, 122.96, 115.50, 115.36, 114.57, 110.15, 109.20, 108.41, 98.12, 60.34, 43.51, 41.69, 41.01, 40.17, 39.89, 26.47. **HRMS**: *m/z*: calcd for C_35_H_35_ClFN_6_O_4_: 657.2387; found 657.2384 [M + H]^+^.

### GNF179-NBD

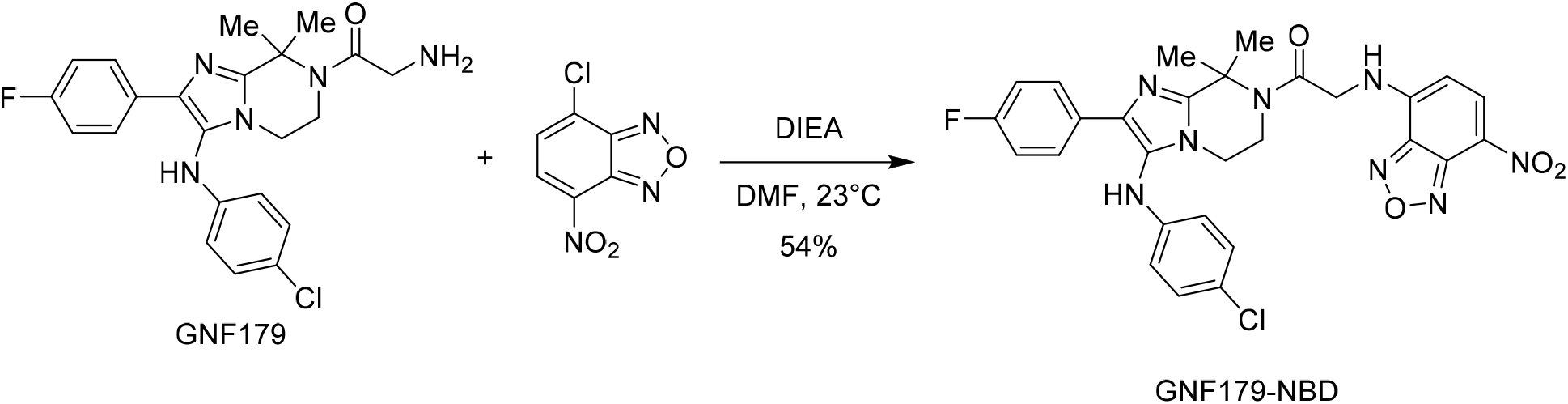

#### GNF179-NBD

To a stirred solution of GNF179 (6 mg, 0.014 mmol, 1.0 equiv.) and 4-chloro-7-nitrobenzofuran (3 mg, 0.014 mmol, 1.0 equiv.) in dry DMF (0.2 mL) was added neat DIEA (4 μL, 0.021 mmol, 1.5 equiv.). The solution was stirred for 16 hours at 23 °C. The reaction was then diluted with EtOAc (2.5 mL) and water (1.2 mL) and the organic layer was collected. The mixture was extracted with additional EtOAc (2 mL). The EtOAc layers were dried over sodium sulfate, filtered, and concentrated. The crude product was purified by silica column chromatography, eluting with Hexanes: EtOAc (40: 60) afforded the title compound (4 mg, 6.77 mmol, 52%) as light brown solid.

#### NMR results

**R**_**f**_ = 0.6 (silica gel, 30:70 Hexanes: EtOAc); ^**1**^H NMR (600 MHz, CDCl_3_) δ 8.35 (d, *J* = 8.4 Hz, 1H), 7.72 – 7.65 (m, 2H), 7.59 (s, 1H), 7.13 (d, *J* = 8.2 Hz, 2H), 6.94 (t, *J* = 8.3 Hz, 2H), 6.57 (d, *J* = 8.2 Hz, 2H), 6.13 (s, 1H), 5.60 (s, 1H), 4.30 (s, 2H), 3.91 (m, 2H), 3.80 (m, 2H), 2.00 (s, 6H). ^**13**^C NMR (151 MHz, CDCl_3_) δ 162.72, 161.08, 147.27, 144.22, 133.82, 129.81, 129.65, 129.64, 127.67, 127.62, 124.62, 122.93, 115.53, 115.38, 114.59, 60.93, 56.09, 41.95, 41.03, 29.80, 26.52. **HRMS**: *m/z*: calcd for C_28_H_25_ClFN_8_O_4_: 591.1666; found 591.1662 [M + H]^+^.

The potency of both GNF179 fluorescent conjugate was assessed via a 72-hour SYBR Green I fluorescence IC_50_ assay.^74^

### Sphingolipid Quantitation

Parasites were grown in culture media containing both human serum (5%) and albumax (0.25%) at 3% hematocrit. Asynchronous parasites (containing ~2/3^rd^ rings) were treated with GNF179 for 4 hours at 5X wild type IC_50_ (25nM). Parasitized cells were then washed with PBS (+ drug for GNF179-treated samples). Samples were then lysed with 5 volumes of 0.15% saponin (in cold PBS) on ice for 20min, and centrifuged and washed extensively. Parasite pellets were flash frozen in liquid nitrogen and stored at −80°C until use.

Protein concentrations were measured using the Bradford Protein Assay for biological (n=3) and technical (n=2) replicates for WT and mutant parasites ± drug. Samples were solubilized using cell lysis buffer comprising 1M Tris, 20% SDS, 0.5M EDTA, β-mercaptoethanol and double distilled water then boiled at 95°C for 5min. Bovine serum albumin (BSA) standards and parasite samples were incubated with Bradford Reagent (BioRad) at room temperature for 15-30min then absorbance read at 595nm in order to derive protein concentrations.

For lipid extraction, parasite samples and controls for lipid analysis were prepared in a 96 well plate comprising a final volume of 50μl per well. To each well, 50μl internal standards comprising ceramide (C12) and sphingomyelin (SM12) were added. Lipids were extracted from parasite and control samples using 900μl dichloromethane/methanol (1:1 v/v). The plate was vortexed overnight at room temperature then centrifuged to pellet out all insoluble material. The supernatant containing lipids was transferred into a new plate and samples subjected to mass spectrometry using a triple quadrupole LC-MS system. Briefly, 7μl of sample was injected into the HPLC running through a C18 Poroshell column into Agilent QQQ 6430 MS/MS. HPLC was optimized under the following conditions: mobile phase A (water/ methanol/ chloroform/ 0.1% formic acid) and mobile phase B (methanol/ acetonitrile/ chloroform/ 0.1% formic acid). Analysis was done on Agilent Quantitative Analysis software and normalized to total protein in the samples.

All sample data sets were analyzed using Graph Pad Prism 4.0. Tables 2.1-2.19 below, show the output from Prism comparing sphingolipid content across all possible combinations of data in treated (+) and untreated (-) wild type (Dd2) and *pfcarl* mutant (KAD452-R3) parasites. Sphingolipids were normalized to protein and reported with respect to the mean ± SEM for n=3 independent experiments. Asterisks included in the tabulated data indicate statistical significance at the 95% confidence interval determined using a two-tailed t-test (* p < 0.05, ** p < 0.01, *** p < 0.001). n.d. denotes samples with insufficient data or lipid contents below the detection limit.

### Immunoblotting to demonstrate Protein Trafficking Defects

*P. falciparum* parasites expressing PfEMP3-GFP or PfSERA5ss-GFP were generated as previously described^41^, with the indicated reporters cloned into pGLUX.1 and their expression driven by the PfCRT promoter*. P. falciparum* trophozoites expressing PfEMP3-GFP or PfSERA5ss-GFP were magnetically-purified (Miltenyi Biotech), incubated with 10µM GNF179 or WEHI-842 for 3hrs at 37°C, and treated with 0.09% saponin-containing inhibitor. Washed pellets were solubilized in 2x Laemmli buffer (Bio-Rad #1610737) then boiled for 3 min. Proteins were then separated by SDS-PAGE, transferred to nitrocellulose and blocked in 1% skim milk. Membranes were probed with mouse anti-GFP (Roche, cat. no. 11814460001) (1:1,000) (primary validation is provided on the manufacturer’s website) or rabbit anti-HSP70 (1:4,000), subsequently followed by probing with species-matched horseradish peroxidase– conjugated secondary antibodies (Cell Signaling Technology, cat. nos. 7074 and 7077) and visualization with enhanced chemiluminescence (Amersham).

For endogenous proteins, 3D7 wild-type *Plasmodium* parasites were treated with the indicated concentrations of GNF179 for 3hr at 37°C, and treated with 0.09% saponin-containing inhibitor. Immunoblotting was performed as described above using antibodies for PfIESP2, PfSERA5 and PfPTP2^75^.

### Sec62 knockdown assay

3D7 wild type and PfSec62-HA-glmS transgenic parasites^45^ were synchronized using sorbitol at ring stage and incubated with 0 mM and 1 mM Glucosamine (Sigma). GNF179, chloroquine and mefloquine were added at increasing concentrations and all cultures incubated for 48 hrs until the late ring stage of the successive cycle. Parasitaemia at each drug concentration was assessed by flow cytometry and all data tabulated using GraphPad Prism.

### Measuring Parasite Translation using ^35^S-Incorporation

The effect of drug treatment on parasite translation was evaluated by quantifying the incorporation of ^35^S-labeled amino acids into newly synthesized protein by adapting a published protocol ^76^. Briefly, synchronized, trophozoite-stage parasites (5% parasitemia, 26-30 hpi) were first washed in methionine-free media three times prior to drug treatment. Parasites were then incubated in various compound concentrations in 24-well plates for 1 hour at 37°C, at a final hematocrit of 5%, and a final concentration of 125 µMci/mL of EasyTag™ EXPRESS^35^S Protein Labeling Mix (Perkin Elmer, USA). Concentrations used for incubation correspond to 100-, 10-, 1-, 0.1- and 0.01-times the IC50 values of KAF156 (15 nM), chloroquine (85 nM) and cycloheximide (750 nM). After incubation, parasites were washed with 1x PBS and then lysed using ice-cold 0.15% saponin in 1x PBS for 20 minutes. All subsequent steps were performed on ice. The saponin pellet was washed with 1x PBS, and then resuspended in 0.02% sodium deoxycholate and supplemented with an equal volume of 16% trichloroacetic acid (TCA) to make a final concentration of 8% TCA. The suspensions were then incubated for 20 minutes on ice before vacuum filtration. To collect the radiolabeled, precipitated proteins, the samples were dispensed onto 0.7µM glass fiber filter discs (Millipore, USA) that had been presoaked in 8% TCA. The vacuum-filtered precipitates were then washed twice with 8% TCA and then, finally, with 90% acetone. The filter discs were allowed to air-dry for at least two hours, transferred into scintillation vials and resuspended in scintillation cocktail (Perkin Elmer, USA). ^35^S counts were obtained for 1 minute using a Beckman Coulter LS 6500 Multi-purpose Scintillation Counter. Counts were normalized to data obtained from untreated parasites.

### New Permeation Pathways Assessment of Sorbitol Synchronization Efficacy after GNF179 treatment

Parasites were cultured as above. Three parasite strains were used, wild-type Dd2, KAD452-R3 and Dd2 ACT S242* (Dd2 *act* -PF3D7_1036800 – with S242* SNV ^21^). Each parasite clone was split into 3 sets of matched cultures. Each set of matched cultures was either treated for 4hr at 37°C with compound (50nM GNF179, 500nM Chloroquine or 5µM Brefeldin A (Sigma Aldrich #B6542)) or an equal volume of DMSO. After drug treatment, cultures were synchronized by treatment with 5% sorbitol (Sigma Aldrich # 240850) for 10min at 37°C, washed once with media and returned to normal culture conditions. 24 hours after treatment, cultures were assessed by thin blood smear and GIEMSA staining for parasite lifecycle stage. The proportion of early (‘ring’) stage parasites to the total parasite proportion was recorded.

### Microscopy Methods Parasite culture

*P. falciparum* asexual blood-stage parasites were cultured in human erythrocytes (3% hematocrit) and RPMI-1640 media supplemented with 2mM L-glutamine, 50mg/L hypoxanthine, 25mM HEPES, 0.225% NaHCO_3_, 10mg/L gentamycin, and 0.5% (w/v) Albumax II (Invitrogen). Parasites were maintained at 37ºC in 5% O_2_, 5% CO_2_, and 90% N_2_. Cultures were stained with Giemsa, monitored by blood smears fixed in methanol, and viewed by light microscopy.

### Export block assay

The KAHRP69-GFP reporter line was generated by amplifying the first 207 bp of the KAHRP gene (PF3D7_0202000), which includes the signal peptide and PEXEL motif, followed by a glycine linker, and cloning upstream of GFP in the pDC2-crt-attP-BSD vector under the control of the *pfcrt* 5’ UTR. The plasmid was integrated by attB x attP recombinase-mediated integration into the Dd2attB parasite line^77^. Dd2attB KAHRP69-GFP ring stage parasites were incubated with either 5×IC_50_ GNF179 (25nM), Brefeldin A (5µg/ml) (Sigma Aldrich) or DMSO-mock treated 16hr prior to imaging. For live-cell imaging, 5 µl of resuspended culture was added to 30 µl of RPMI-1640 media (without Albumax II) containing Hoechst 33342 (1µg/ml) (Sigma) and imaged at room temperature after 5min incubation at 37°C. For experiments with ER-Tracker^TM^ Red (Thermo Fisher), the dye was added to a final concentration of 1µM 30min prior to imaging. Indirect Immunofluorescence assays (IFAs) were performed in suspension. Cells were fixed in 4% (v/v) formaldehyde (Thermo Fisher Scientific) for 1h at RT followed by a second fixation step supplementing the 4% formaldehyde solution with 1mM Cysteine and CaCl_2_ and subsequent incubation over night at 4°C. The cells were permeabilized on ice using 0.05% Triton X-100 in 1×PBS for 5min and autofluorescence was quenched using 50 mM glycine for 10 min. After two washes in 1×PBS the cells were resuspended in 1% (w/v) bovine serum albumin (BSA) in 1×PBS blocking buffer and incubated with the appropriate dilution for each primary antibody used (1/200 for anti-ERD2, 1/200 for anti-PDI (Mouse anti-PDI (1D3), Enzo Life Sciences, Cat. No. ADI-SPA-891-D), 1/500 for anti-GFP) followed by an incubation with the species-specific corresponding secondary antibody (Alexa Fluor 488-, 594- or 647-conjugated goat anti mouse or rabbit antibodies, Thermo Fisher) diluted 1:2000 in 1% BSA in 1×PBS.

Parasites were imaged using a Nikon Eclipse Ti-E wide-field microscope equipped with a sCMOS camera (Andor) and a Plan-apochromate oil immersion objective with 100× magnification (1.4 numerical aperture). A minimum of 27 Z-stacks (0.2 μm step size) were taken of each parasitized RBC. NIS-Elements imaging software (Version 5.02, Nikon) was used to control the microscope and camera as well as to deconvolve the images (using 25 iterations of the Richardson-Lucy algorithm for each image) and perform 3D reconstructions. ImageJ (Fiji) (version 2.0.0-rc-68/1.52h) was used to crop the images, adjust brightness and intensity, overlay channels and prepare montages.

## Supporting information

Table S2. S. cerevisiae mutations detected by whole genome sequencing

## Author Contributions

Parasite functional assays were conceived by G.L., D.M., D.A.F., A.C. and E.AW. Parasite functional assays, including gene knockdowns, drug synergy experiments and conditional knockdowns, were performed by G.L., D.M, J.K.T. and BY.Z.. Parasite *in vivo* transation assays were performed by F.R. *S. cerevisiae* resistance evolutions and functional assays were performed by E.V., J.Y., A.L.C., P.K. G.G. and S.O. Whole-genome sequencing was performed by the UCSD Institute for Genomic Medicine Core Facility and analyzed by M.L. Fluorescent compounds were synthesized by D.S, T.J. and J.H. E.A.W. wrote the manuscript. All authors read and approved the manuscript. The authors declare no conflicts of interest or competing financial interests.

## Acknowledgements

The authors would like to thanks the members of the Winzeler lab for technical support, reagents and critical reading of the manuscript. GL is supported by an AP Giannini post-doctoral fellowship. E.A.W. is supported by grants from the NIH (5R01AI090141 and R01AI103058) and by grants from the Bill & Melinda Gates Foundation (OPP1086217, OPP1141300) as well as by Medicines for Malaria Venture (MMV). DAF gratefully acknowledges funding from the Medicines for Malaria Venture and the Bill & Melinda Gates Foundation. M.L. was supported in part by a Ruth L. Kirschstein Institutional National Research Award from the National Institute for General Medical Sciences, T32 GM008666.

## Data Availability

All genome sequences for the 13 IZP-resistant *S. cerevisiae* strains have been placed in the short-read sequence archive (http://www.ncbi.nlm.nih.gov/sra) under accession code STUDY: PRJNA381796 (SRP107357). Whole-genome sequences for KAD452-R3 can be downloaded from (located on NAS server: Victoria, collaborative sequencing projects, Lamonte_KAF156R)

## Supplemental Figures and Tables

**Figure S1.**
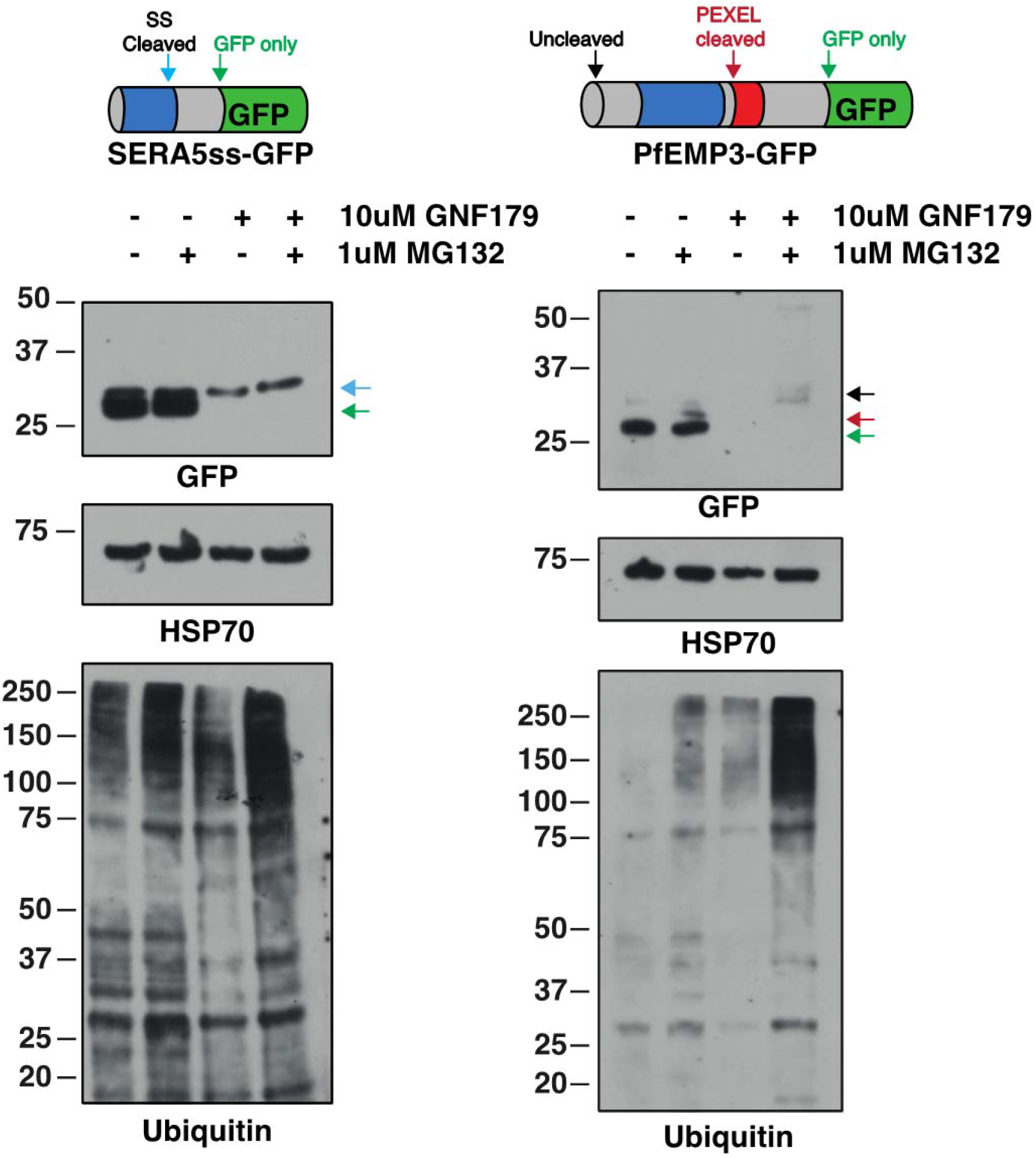
does not induce a significant ubiquitin response in *P. falciparum*. Western blots of *P. falciparum* strains expressing SERA5ss-GFP or PfEMP3-GFP treated with 10uM GNF179 and/or 10µM MG132 for 3 hours were probed with anti-GFP, anti-HSP70 and anti-Ubiquitin antibodies. For the secreted SERA5ss-GFP reporter, the blue arrow indicates the signal-sequence cleaved form and the green arrow indicates the GFP-only species. For the exported PfEMP3-GFP reporter, the black, red and green arrows indicate the full-length, PEXEL-cleaved and GFP only species respectively.

**Figure S2.**
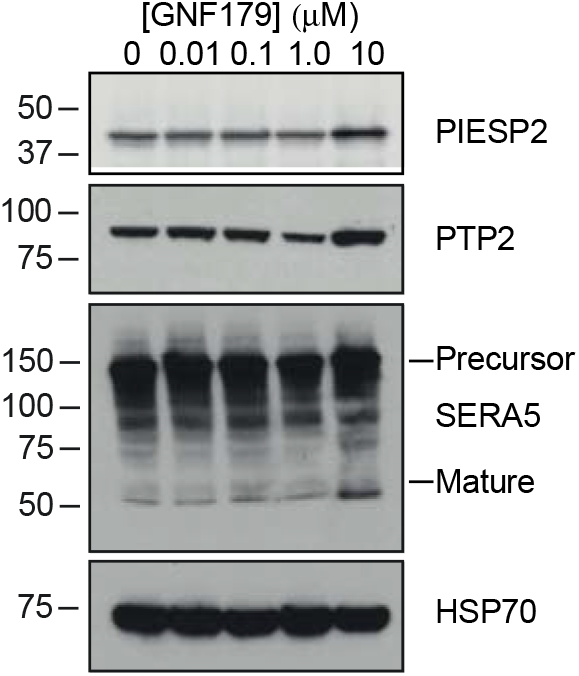
Western blot showing effects by a 3-hour pulse of increasing concentrations of GNF179 on known ER-trafficked proteins in P. falciparum. PIESP2 and PTP2 are examples of PEXEL-containing exported proteins. SERA5 is a secreted protein while HSP70 is a cytosolically expressed protein that lacks a signal sequence required for ER entry.

**Figure S3.**
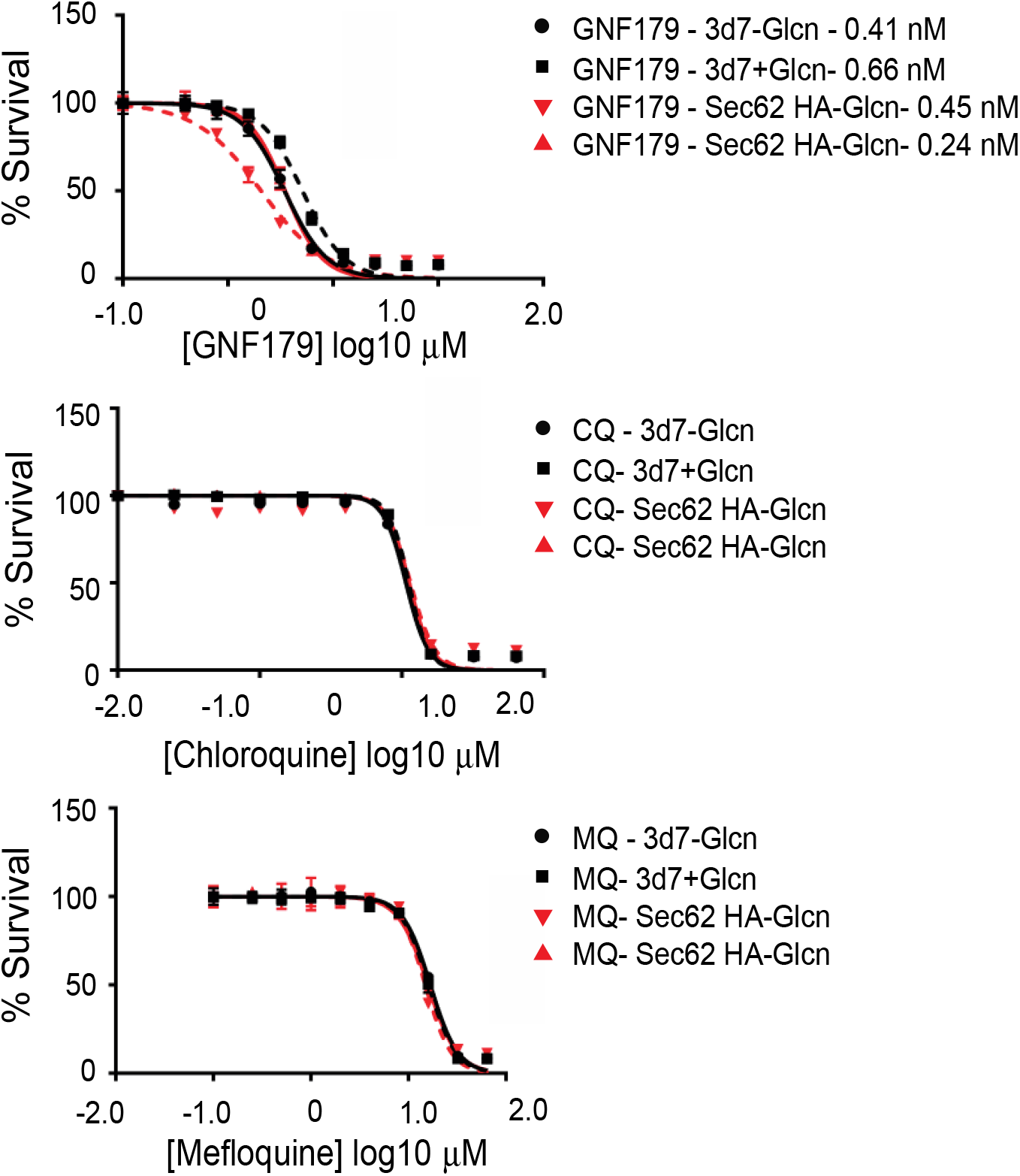
Dose response curves for GNF179 in the SEC62 knockdown parasites, compared to wildtype 3D7 parasites, with and without the addition of N-acetyl Glucosamine (Glcn) for the 3 compounds indicated, GNF179, Chloroquine (CQ) and Mefloquine (MQ). Values are for three independent replicates.

**Figure S4.**
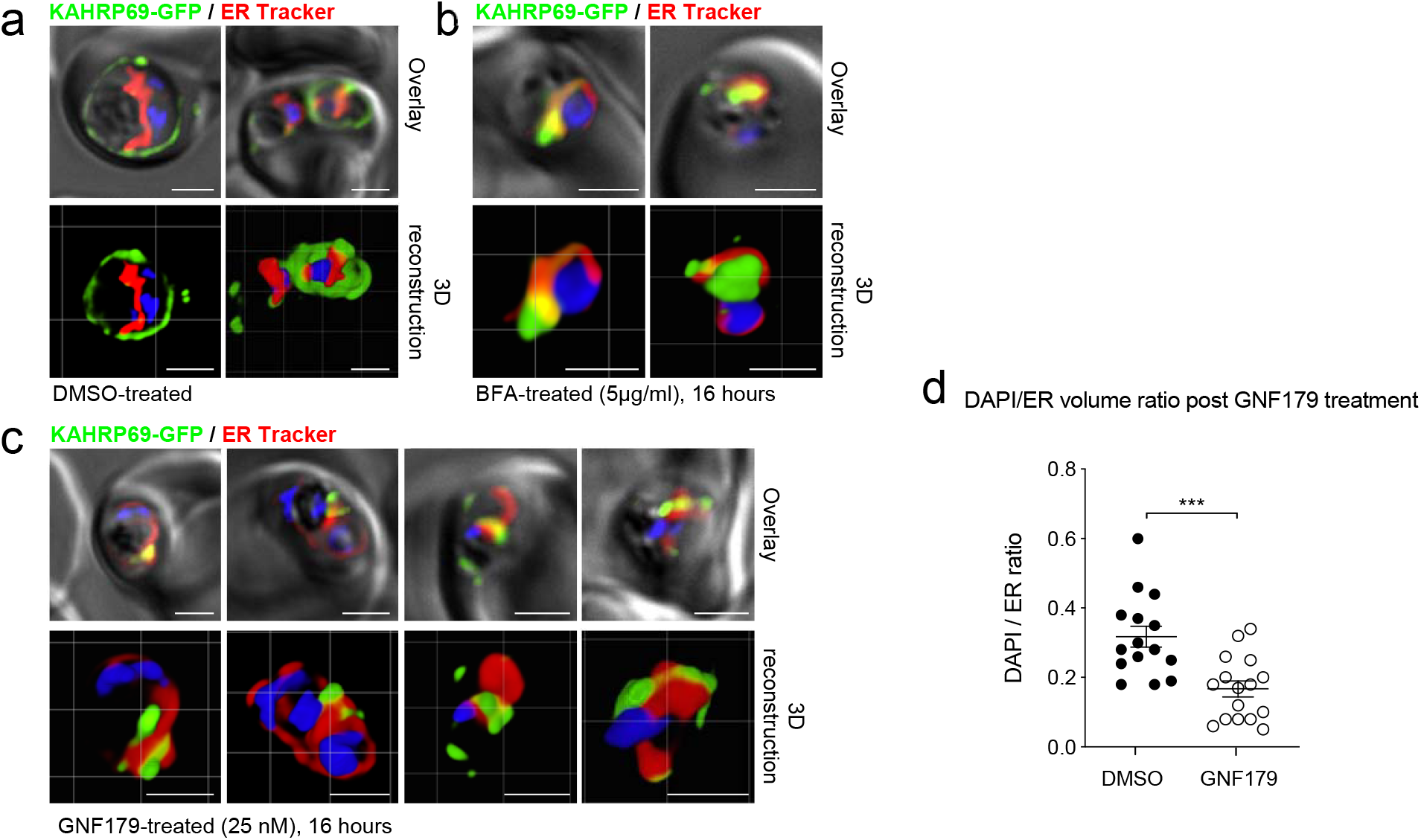
GNF179 treatment increases volume of ER relative to nucleus in Dd2attB KAHRP69-GFP parasites. Red, ER Tracker; Green, GFP; Blue, DAPI. (a) DMSO-treated parasites. (b) Treatment with 5 µg/ml Brefeldin A for 16 hours. (c) Treatment with GNF179 for 16 hours. (d) Quantitation of the 3D Volume ratio of DAPI to ER tracker in 12 images. Statistical significance was determined using a paired, two-tailed t-test. Scale bars: 2 μm

**Figure S5.**
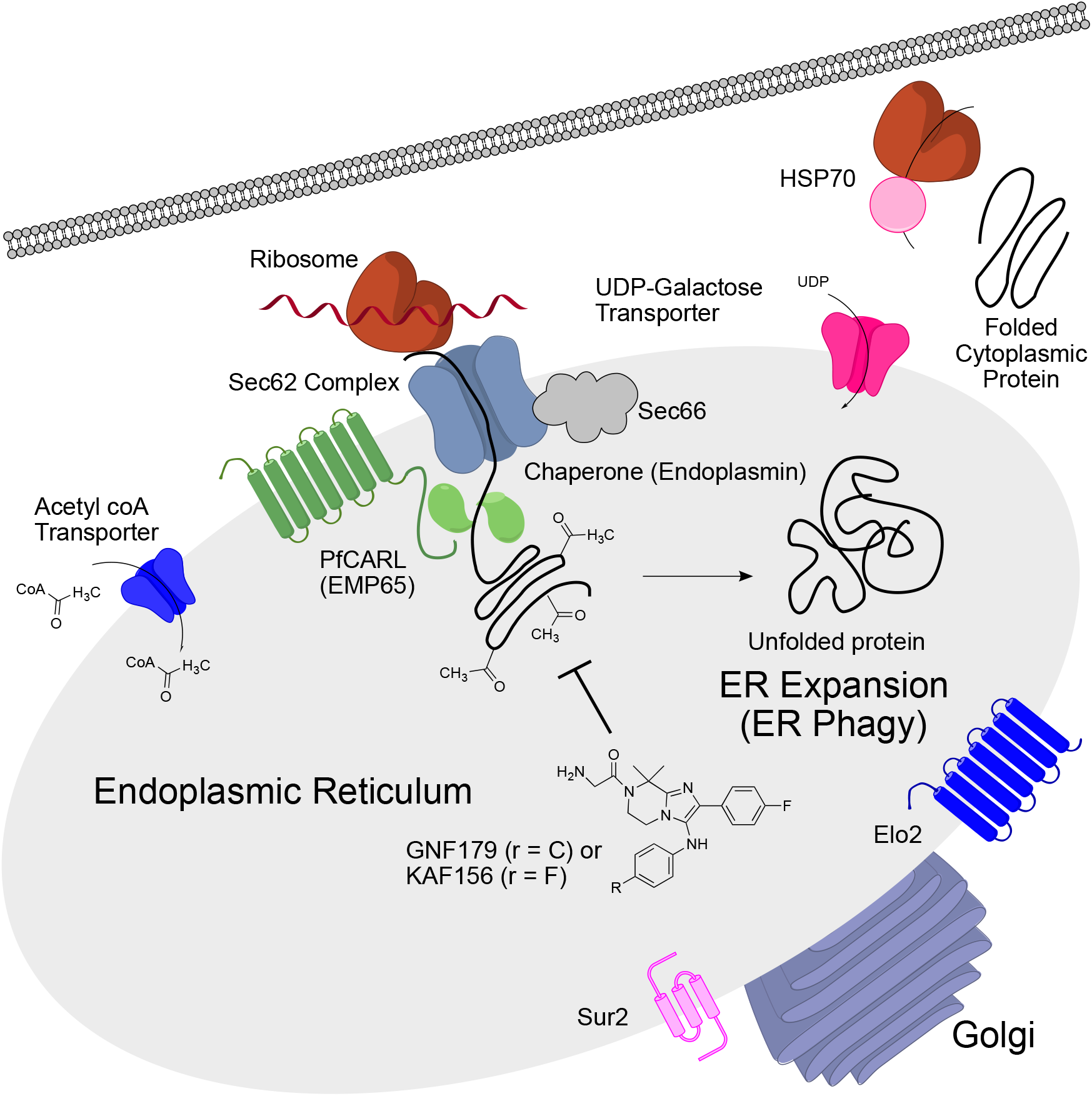
Model that shows how IZP treatment could produce the sets of cellular phenotypes that are observed after treatment. In the model, IZPs inhibit the production of properly folded proteins, potentially by interfering with post-translational modification or folding. Mutations that slow the process of protein processing, as those in the Acetyl Coa transporter, *pfcarl* or *sec66* are beneficial. The accumulation of unprocessed proteins leads to ER expansion. Mutations in the autophagy and sphingolipid pathways (*elo2*, *sur2*) in yeast change the balance of ER phagy, potentially also increasing transit time and providing a slight growth advantage in the presence of GNF179 or KAF156.

**Table S1.** Sequencing and alignment statistics for the 13 GNF179-resistant *S. cerevisiae* strains sequenced in this study. All genome sequences for the 13 IZP-resistant *S. cerevisiae* strains have been placed in the short-read sequence archive (http://www.ncbi.nlm.nih.gov/sra) under accession code STUDY: PRJNA381796 (SRP107357).

| Clone Name | Total Base Pairs Sequenced | Mean Coverage (X) | Percent Reads Aligned to Reference | Percent Bases with 20 or More Reads | IC50 (uM) |
| --- | --- | --- | --- | --- | --- |
| **Green-Monster** | 557,641,556 | 42.3 | 98.7 | 95.0 | 45.39 ± 5.7 |
| **GNF179-G72** | 1,085,965,416 | 59.2 | 99.2 | 94.0 | 63.45 ± 2.6 |
| **GNF179-R1-2** | 483,257,249 | 31.1 | 99.1 | 83.7 | 80.12 ± 14.6 |
| **GNF179-R10-2** | 868,810,757 | 49.1 | 99.4 | 93.4 | 90.31 ± 14.8 |
| **GNF179-R12-2** | 1,238,282,174 | 71.8 | 99.4 | 95.0 | 67.22 ± 10.4 |
| **GNF179-R12h2** | 504,425,192 | 30.9 | 99.4 | 85.3 | 89.65* |
| **GNF179-R13-2** | 938,610,879 | 58.8 | 99.5 | 94.4 | 59.68 ± 11.5 |
| **GNF179-R14-2** | 719,981,126 | 46.4 | 99.7 | 92.0 | 77.83 ± 7.3 |
| **GNF179-R18g1** | 510,038,211 | 30.7 | 99.5 | 85.2 | 137.66 ± 5.1 |
| **GNF179-R19g2** | 607,361,091 | 35.3 | 99.4 | 89.1 | 109.35 ± 23.1 |
| **GNF179-R7-2** | 1,137,607,696 | 66.7 | 99.4 | 94.6 | 71.11 ± 9.5 |
| **GNF179-R8h2** | 654,463,110 | 38.2 | 99.3 | 90.0 | 138.05* |
| **GNF179-R9-2** | 945,657,737 | 61.3 | 99.4 | 94.2 | 81.18 ± 6.3 |
| **GNF179-R9f2** | 697,615,853 | 40.2 | 99.4 | 87.0 | 94.7 ± 41.3 |

\*Replicate not available

**Table S2. Complete set of mutations observed in *S. cerevisiae in vitro* resistance evolutions. (Attached Excel Table)** Clone #: The numerical designation for each *S. cerevisiae* GNF179 resistant clone in this study. Chr #: Chromosome # which harbors the identified mutations.

**Table S3.**
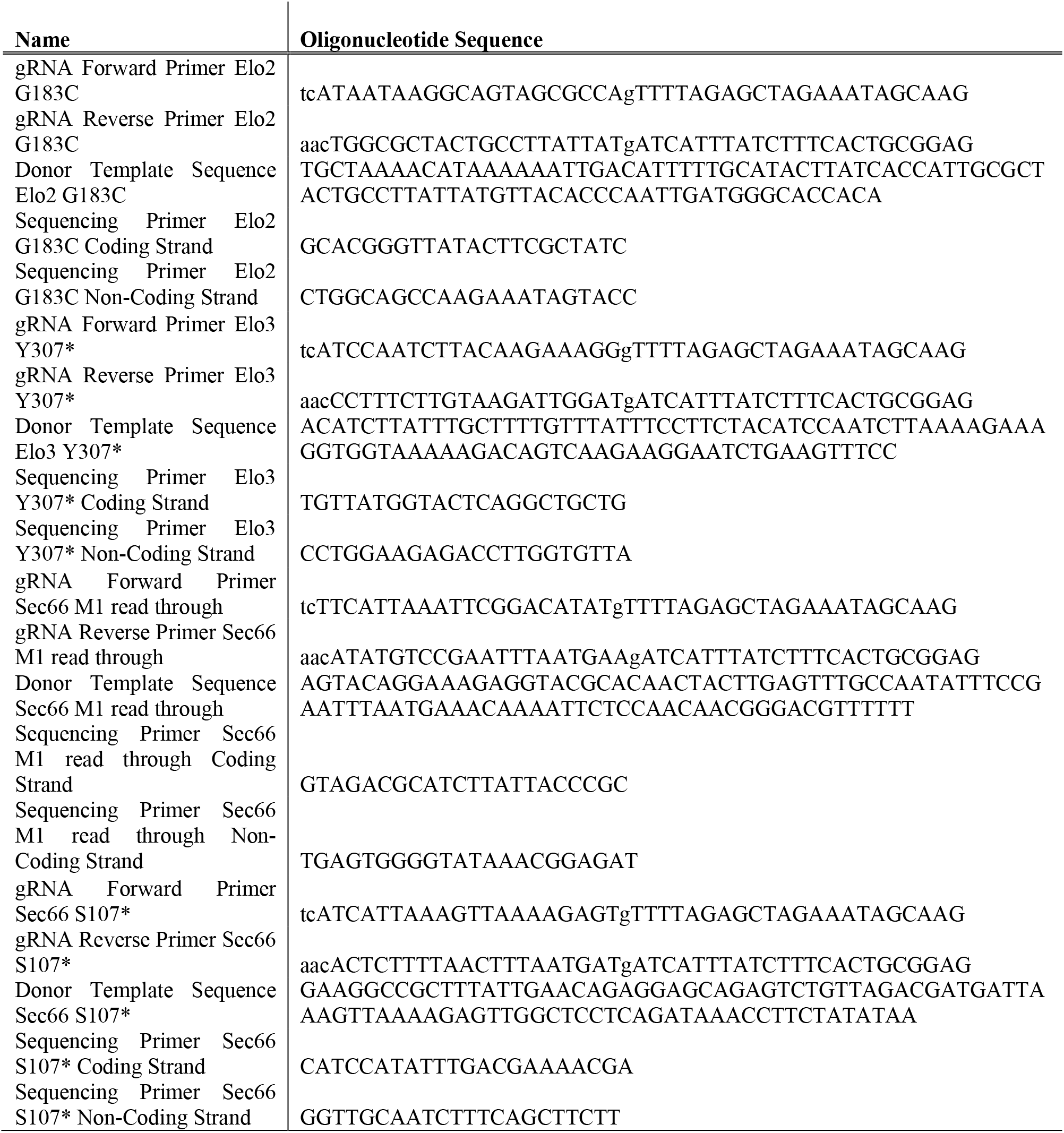
Oligonucleotides used in this study.

